# Standard operating procedure for somatic variant refinement of tumor sequencing data

**DOI:** 10.1101/266262

**Authors:** Erica K. Barnell, Peter Ronning, Katie M. Campbell, Kilannin Krysiak, Benjamin J. Ainscough, Cody Ramirez, Nick Spies, Jason Kunisaki, Jasreet Hundal, Zachary L. Skidmore, Felicia Gomez, Lee Trani, Matthew Matlock, Alex H. Wagner, S. Joshua Swamidass, Malachi Griffith, Obi L. Griffith

## Abstract

**Purpose:** Manual review of aligned sequencing reads is required to develop a high-quality list of somatic variants from massively parallel sequencing data (MPS). Despite widespread use in analyzing MPS data, there has been little attempt to describe methods for manual review, resulting in high inter- and intra-lab variability in somatic variant detection and characterization of tumors.

**Methods:** Open source software was used to develop an optimal method for manual review setup. We also developed a systemic approach to visually inspect each variant during manual review.

**Results:** We present a standard operating procedures for somatic variant refinement for use by manual reviewers. The approach is enhanced through representative examples of 4 different manual review categories that indicate a reviewer’s confidence in the somatic variant call and 19 annotation tags that contextualize commonly observed sequencing patterns during manual review. Representative examples provide detailed instructions on how to classify variants during manual review to rectify lack of confidence in automated somatic variant detection.

**Conclusion:** Standardization of somatic variant refinement through systematization of manual review will improve the consistency and reproducibility of identifying true somatic variants after automated variant calling.

## INTRODUCTION

Identification of high-quality somatic variants from aligned sequencing data first requires automated processing of aligned sequencing reads by somatic variant callers such as: EBCall,^1^ Mutect,^2^ SomaticSniper,^3^ Strelka,^4^ VarScan2,^5^ and Virmid.^6^ Currently, there is no community consensus as to the single, most appropriate variant caller.^7,8^ A resulting common practice is to utilize the union or intersection of multiple variant callers to develop a preliminary list of potential somatic variants with the hope of a specific capture of all legitimate mutations.^9^ This approach typically requires subsequent refinement of true somatic variants through the elimination of false positives and confirmation of true positives to develop a final list of reputed somatic variants.

Despite variant callers capability to account for zygosities, ploidies, and sequencing artifacts, additional filtering and refinement of somatic variants is often required to correct for differences in data generation methods, sample differences, and variant caller inaccuracies and idiosyncrasies. Typically, this involves manual inspection of aligned reads (i.e. manual review) to eliminate false positives and confirm true positives. Manual review allows trained individuals to incorporate information that is neglected, poorly modeled, or unavailable to automated variant detection to improve somatic variant refinement. For example, the human eye can discern misclassifications attributable to overlapping errors at the ends of sequencing reads, preferential amplification of smaller fragments, poor alignment in areas of low complexity, and combinations of these, and other factors.^10^ Due to the computational inability to synthesize these complex patterns, automated methods to refine somatic variants identified by somatic variant callers have not yet replaced manual review in many workflows.

Manual inspection of somatic variants identified by automated somatic variant callers is an important aspect of the sequencing analysis pipeline and is currently the gold standard for variant refinement.^11,12^ Additionally, manual review is prevalent in the clinical diagnostic and molecular pathology setting, and has been cited in many clinical guidelines for NGS-based diagnostics.^13–15^ However, despite its extensive use, somatic variant refinement strategies are often unstated or only briefly mentioned in articles that practice post-processing of automated variant calls.^11,16–19^ Lack of formalized procedures for somatic refinement allows for high levels of inter- and intra-lab variability and can prevent reproducibility of results. Thus, development of a procedure to standardize and systematize somatic variant refinement would improve the overall quality of sequencing pipelines.

Here we present a standard operating procedure (SOP) for manual review to help standardize somatic variant refinement. We first detail instructions for downloading and using the previously described and publicly available Integrative Genomics Viewer (IGV)^11,12^ software to properly visualize somatic variants during manual review. Additionally, we describe the IGVNavigator (IGVNav), software which facilitates manual review via a graphical user interface. This interface allows for categorization of calls into four distinct classes with the added ability to tag variants with 19 common patterns observed during manual review. Adoption of a standard method for somatic variant refinement through this manual review SOP will streamline somatic variant refinement and systemize input and output reports for downstream analysis.

## MATERIALS AND METHODS

### Manual Review using Integrative Genomics Viewer (IGV)

The Integrative Genomics Viewer (IGV)^11,12^ is a high-performance visualization tool for analysis of large genomics datasets. It supports both array-based and next generation sequence data and provides numerous genomic annotation features. Although IGV has many applications, this standard operating procedure (SOP) will review IGV components that can be used to conduct somatic variant refinement through manual review of variants identified by somatic variant callers.^11,20^ While we have chosen IGV to develop our manual review SOP and demonstrate key concepts, a number of other genomics viewers are also available including: Savant,^21^ Trackster,^21,22^ and BamView.^23^ Many of the following concepts (apart from IGV-specific instructions), should be applicable to most other genomic viewers.

The IGV Desktop application can be downloaded at http://software.broadinstitute.org/software/igv/. This site provides an overview of the development and functions of IGV as well as instructions for software download. Download instructions are available at http://software.broadinstitute.org/software/igv/download. IGV is available for all major operating systems. The presented SOP is consistent with IGV version 2.4.8 using macOS High Sierra Version 10.13.3.

### Information on Integrative Genomic Viewer (IGV) features

The IGV User Guide can be accessed online (http://software.broadinstitute.org/software/igv/UserGuide). This user guide provides detailed information on the interface, navigating through data, loading data, etc. Briefly, the IGV application view is broken down into three parts: 1) the Genome Ruler, 2) Data Tracks, and 3) the Genome Features (Figure 1). The Genome Ruler provides navigation features to properly center genomic loci of interest. For example, the reference genome dropdown menu allows you to switch the current genome being visualized (default is human b37); the variant coordinates show the current field of view; the popup text behavior button allows you to change the text display in data panels; and the zoom buttons allow you to expand and contract the data tracks’ field of view. The Data Tracks are loaded individually, whereby each horizontal track represents one experiment or sample. In Figure 1, there are two loaded tracks (a normal track and a tumor track). Each data track consists of a coverage track and read strands. Clicking on, or hovering over the coverage track will show information about the loci of interest. Read strands ideally represent one molecule that was sequenced and aligned to a reference. In default settings, sequenced bases that disagree with the aligned reference sequence are highlighted. In the example provided, the highlighted bases on many tumor read strands support a variant thymine (T) allele in place of the reference guanine (G). The Genome Features provide additional information that can be used to supplement manual review. The reference DNA sequence track and the reference protein sequence track are always loaded using default settings. Additional server tracks that are optionally loaded will be populated in the Genome Features field (Figure 1). There are several additional IGV features are used to identify specific sequencing patterns during manual review, which are outlined in the ‘Helpful Hints’ sections for many examples below.

**Figure 1.**
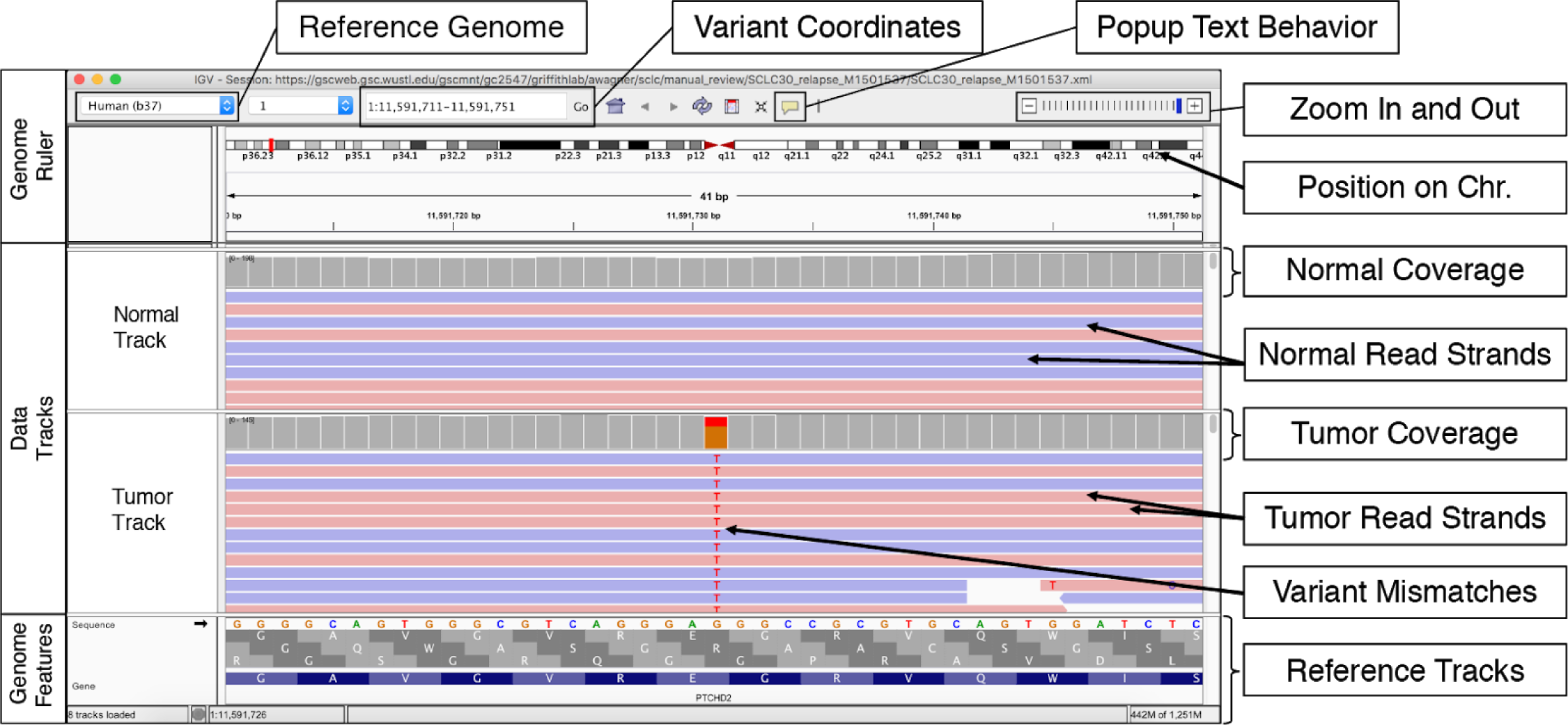
Example of the IGV Interface with associated features relevant to manual review. The IGV interface is divided into three parts. The Genome Ruler section details information about the genome assembly being visualized (Reference Genome), the coordinates currently being visualized (Variant Coordinates), toggle for popup text (Popup Text Behavior), and provides navigation/display controls (Zoom In and Out). In this example, a portion of the human chromosome 1 (build 37) is shown. The central section of IGV displays Data Tracks. In this case, short read DNA alignment data (e.g., bam file) are shown for a normal sample and tumor sample loaded by the user. When alignments are colored by read strand (as recommended) reads are represented as red and blue bars, whereby mismatches with the reference genome are highlighted by base. Coverage tracks summarize the total read depth at each base position. The Genome Features section shows the reference sequence itself, the amino acids for all three possible reading frames, and the gene associated with this locus. The default gene track available with IGV is shown (RefSeq). Many other data formats and sources can be loaded as data tracks or genome features.

### IGV Input Files

The IGV software supports a variety of different input files for genomic sequencing visualization. The “File” drop down on the IGV command bar permits visualization of the various supported input files. Within this dropdown, viable options including: “Load from File”, “Load from URL”, or “Load from Server”, etc. Our lab typically uses the Genome Modeling System^24^ to produce IGV URLs that can be directly uploaded with the “Load from URL” option. The, “Load from Server” option allows one to download tracks from servers that IGV supports such as The Cancer Genome Atlas (TCGA data) or Ensembl, etc. These tracks can assist in the manual review process, for example, the Common SNPs tracks can help ascertain if a variant is a common polymorphism or a true somatic variant. This feature is described in more detail below.

There are at least three different types of sequence experiment designs that require different manual review setup. These include (1) analysis when only a tumor sample is available, (2) analysis when tumor samples and normal samples can be compared, and (3) analysis when you have additional information beyond tumor-normal comparison (e.g. RNA-sequencing data, relapse data, etc.). As mentioned above, it is recommended that all tracks (samples) associated with the same individual should be loaded and analyzed together. Given that some variants might be unique to individual samples, (e.g. a variant was identified in the relapse but not identified in the primary tumor), it might be necessary to merge multiple variant files into one comprehensive file for IGVNavigator input (see below). For reference, we have outlined some caveats associated with each of these three different types of analyses:

#### Tumor Sample Only

If only tumor DNA is available, somatic variants must be assessed by evaluating only the tumor track. Given the difficulty of evaluating true somatic variants with tumor only samples, it may be helpful to load a population SNP track within the IGV session to determine if the variant being evaluated is common in the human population (e.g. 1000 Genomes,^25^ ExAC,^26^ gnomAD^26^). If a variant exists at a very low (e.g. <5%) or zero population frequency within these databases, it increases the likelihood that the variant is somatic. However, *de novo* and less common germline polymorphisms, which could be reliably removed by observation of a matched normal sample, might still pass manual review and other filtering approaches.

#### Tumor Sample + Normal Sample

When tumor DNA and normal DNA are available, they can both be loaded within the IGV session. This increases the ability to label true somatic variants during manual review due to the comparison with the normal track(s) through the elimination of germline variation and systemic artifacts. It is important to note that hematologic tumor types (and possibly other tumor types) may display tumor contamination in the normal sample. Refer to Tumor Normal (TN) variant for information on how to properly assess such variants.

#### Tumor Sample + Other (RNAseq, Relapse, Metastasis, etc) +/− Normal Sample

When tumor DNA, normal DNA and other DNA or RNA are available, they can all be loaded within a single IGV session. Support from multiple tumor tracks increases reviewer confidence that the variant in question is real. Support from RNA sequencing data can be especially convincing and can be used to confirm that somatic mutations are expressed. As mentioned, multiple tracks can be loaded into the same IGV session; however, increasing the total number of tracks can increase the time required for the IGV session to load, especially if sequence files are hosted remotely. Two approaches can be utilized to mitigate load time: 1) use a faster network connection to download remote data locally or 2) use the IGV downsample reads feature to downsample the number of visualized reads. This feature can be implemented using the IGV preferences panel (View → Preferences → Alignments → Downsample reads). When downsampling reads, it is important to consider that there might be an increase in apparent visual artifacts (e.g., variant support in reads in a single direction when in reality, the variant is supported by reads in both directions - see Directions (D) variant for more information). Additionally, downsampling reads can potentially eliminate variants with a low variant allele frequency from view, therefore this feature should be used with caution.

### Manual review is streamlined with IGVNavigator

IGVNavigator (IGVNav) is a tool that expedites manual review of somatic variants. As an input, IGVNav requires a bed (or bed-like) file with variant coordinates and outputs an annotated version of the input file. The variant annotation includes the call (i.e. somatic, germline, ambiguous, or fail), tags to provide additional information about a variant, and a notes section for free text. IGVNav can be downloaded from the Griffith Lab GitHub Repo (https://github.com/griffithlab/igvnav). To install IGVNav for MacOS, the program should be downloaded (IGVNav.zip), unzipped, and added to Applications folder. IGVNav currently only supported on MacOS.

### How to Use IGVNav

After opening the IGV software and loading the desired tracks for review, you can open the IGVNav software. It is important to note that IGV must be open before opening IGVNav. After initiation of the IGVNav software, you will be prompted to open an input file for manual review. The input file for IGVNav is a tab delimited, five column bed (or bed-like) file. The five columns correspond to chromosome, start coordinate, stop coordinate, reference allele, and called variant allele for each SNV/indel. While input file does not require a header, it is recommended to include a header with the following labels: “chr, start, stop, ref, var, call, tags, notes”. For variants that have not yet been manually reviewed, the call, tags, and notes columns should be blank (Figure 2B). The IGVNav interface, and associated features, are shown in Figure 2A. The top of the interface allows one to select if the input file is 1-based, whereby the default option is 0-based. Below this option is the navigation bar, which permits you to navigate through your input variant list. The “S” button next to the navigation bar will sort alignments by base so that mismatched variants will appear at the top of IGV at the centered loci. Below the navigation bar is a section that shows the current variant being visualized and the total number of variants in the input file. To directly navigate to a variant of interest, you can edit this section using the keyboard and select the “Go” button. The three horizontal bars display the coordinate information for the current variant. The first bar details the chromosome, start, and stop position; the second bar shows the reference allele; and the third bar shows the variant allele. This information should reflect the coordinate information listed in the Genome Ruler of the IGV interface. The “Call” section allows the manual reviewer to select each manual review call. The options are mutually exclusive and include: somatic “S”, germline “G”, ambiguous “A”, and fail “F”. The “Tags” section allows the manual reviewer to associate tags to a variant being evaluated. It is important to note that tags can be used for any call (S, G, A, or F), however, they are especially important for ambiguous and failed calls to inform future reviewers what was considered when labeling the variant during manual review. The IGVNav interface also contains a notes section, which allows for free text to reference patterns like dinucleotides, complex variants, adjacent structural variants, etc. At any point during the manual review, the call, tags, and notes selections can be printed to the original input file using the “Save” button (Figure 2C). IGVNav does not automatically save the input file with new data, so be sure to frequently click the save button at the bottom of the IGVNav interface.

**Figure 2.**
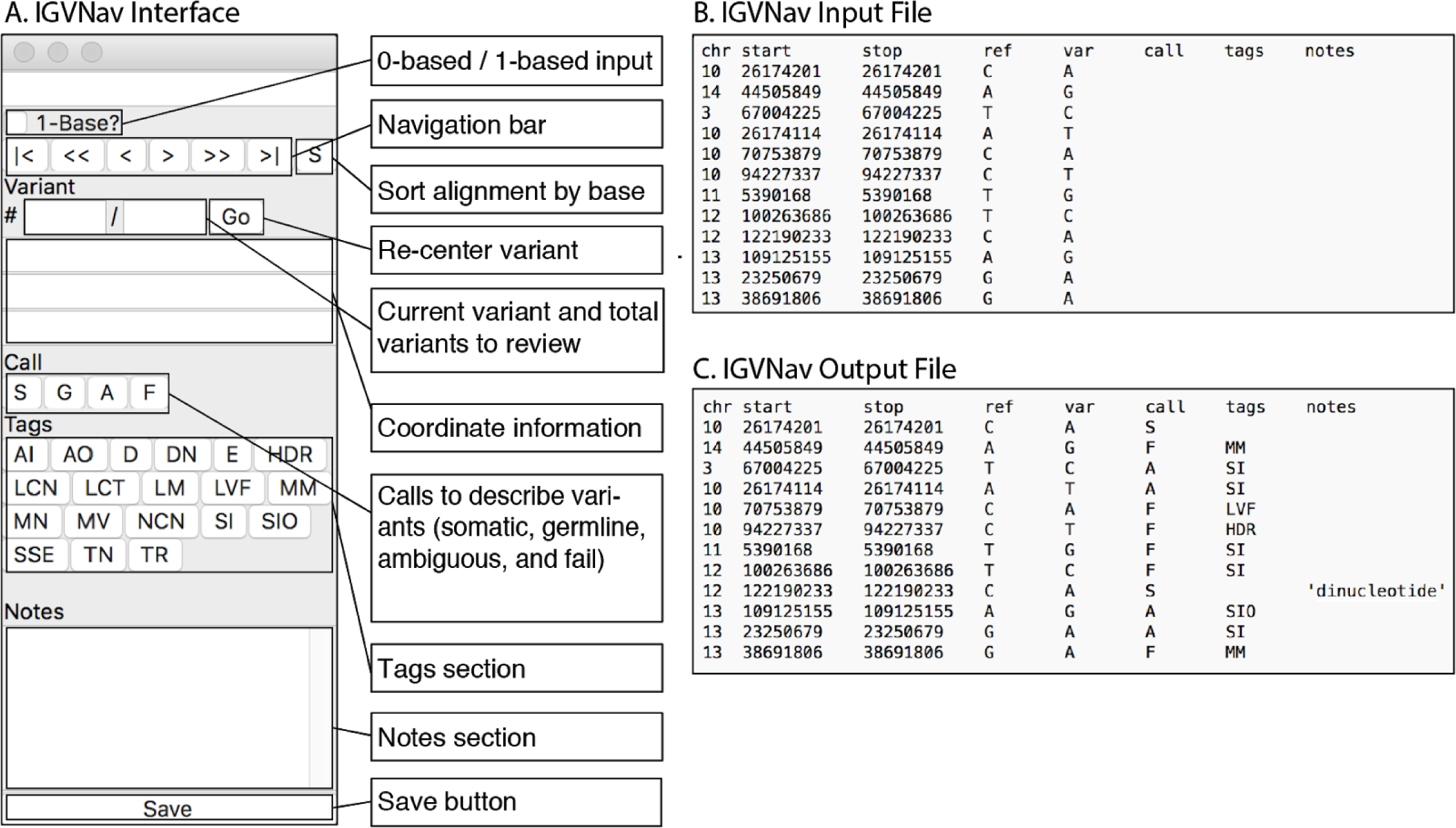
Examples of IGVNav interface and associated features including input and output files. **A.** IGVNav is a simple plugin for IGV that provides a separate application window for recording results of variant manual review. The “1-Base?” button can be selected for 1-base input files (default is 0-base). The “S” button will sort the read strands in the data tracks so that mismatches appear first. The navigation bar allows for movement between variants and displays the variant information. The Call, Tags, and Notes sections allow manual reviewers to annotate variants, which will be reflected in the output file. The save button is used to update the output file. **B.** An IGVNav input file consists of a header line and data for at least the first five columns (chr=chromosome, start, stop, ref=reference base, and var=variant base). Each line represents a variant that will be individually visualized using the Integrative Genomic Viewer (IGV). **C.** During manual review, the input file is updated by clicking on the save button. This will print the call, tags, and notes associated with individual variants to the original input file.

### Step-by-Step guide for setting up manual review

Setting up a manual review session requires following six discrete steps (Figure 3A). First, an IGV session should be opened and the desired reference genome should be selected. Second, the IGV session should be populated with tracks through either a URL or an input file. Step three, which is optional, allows for population of additional tracks that can assist in calling variants. This includes loading annotation tracks like Ensembl or Common SNPs. Step four is also optional, however, we recommend that all tracks be colored by read strand to determine directionality (D) of reads during review. This requires right clicking on each data track and coloring alignments by read strand. After initial set-up of the IGV session, step five requires opening an IGVNav session. After initiating the application, you will be prompted to complete step six, which is to load the tab-delimited .txt or .bed file, whereby the first five columns are populated based on variant information (Figure 2B).

**Figure 3.**
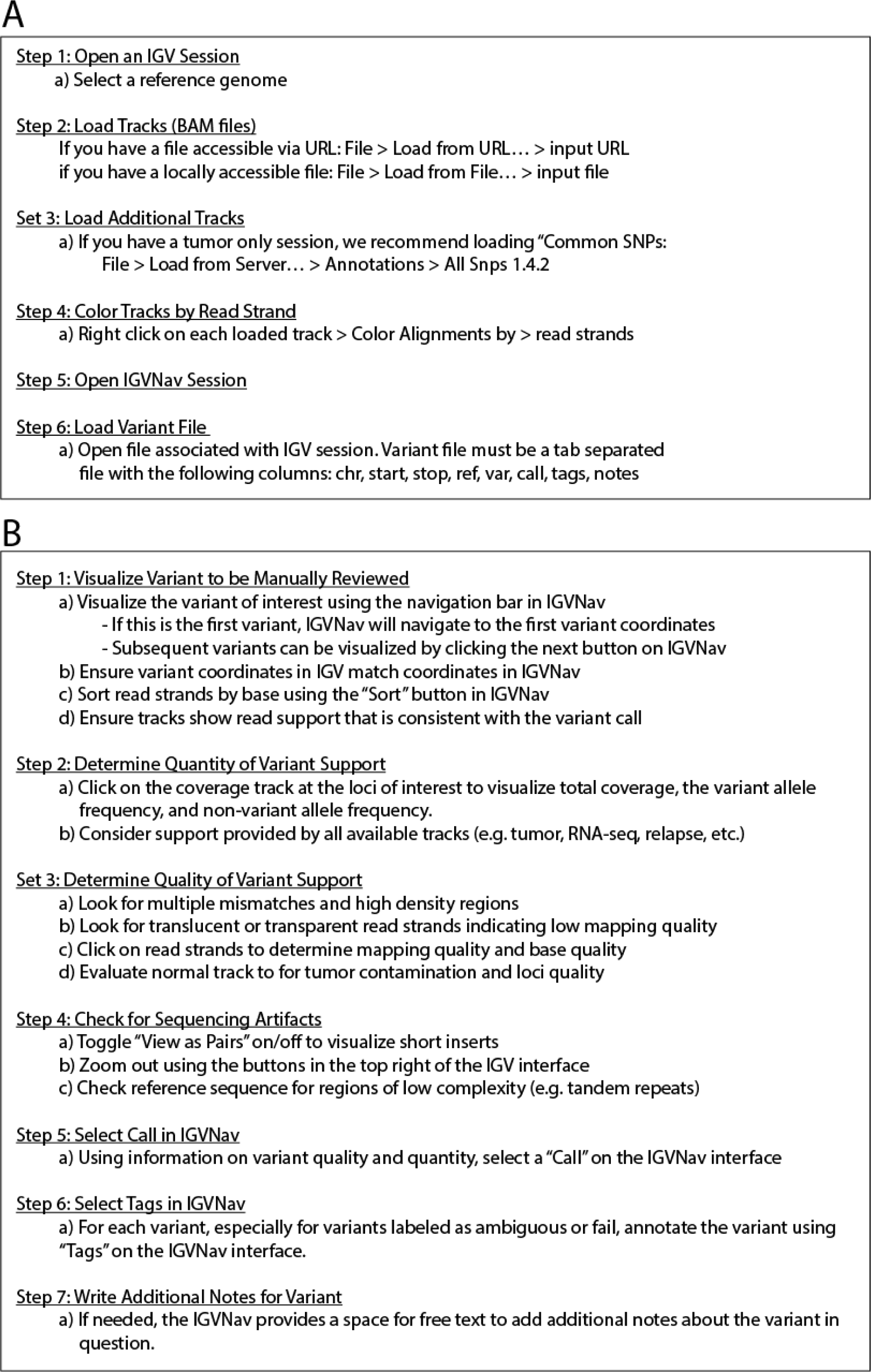
Step-by-Step instructions for setting up and executing manual review of putative somatic SNVs or indels. **A.** Method for setting up IGV_2.4.8 and IGVNav Sessions for manual review. **B.** Method for analyzing each variant during manual review.

### Step-by-Step guide for performing manual review

After initial setup of IGV and IGVNav, there are seven additional steps that must be followed to properly review each variant (Figure 3B). First, you must visualize the variant to be manually reviewed. This can be accomplished by either using the navigation bar in IGVNav or by manually inserting the coordinates into the Genome Ruler section on the IGV interface. During this step, you must also ensure that the coordinates in the IGV Genome Ruler match the IGVNav coordinate information. At this point, it is helpful to sort the read strands by the base to allow for mismatches to be visualized first. This can be accomplished by using the “sort” button on IGVNav or by using the IGV options (right click on track > Sort alignment by > base). Finally, you must ensure that the read support within IGV is consistent with the variant being evaluated. For example, if the variant called is C>A, the centered loci should show a cytosine in the Genome Features reference track and there should be read support for adenine in the tumor tracks. If there is no visualizable variant support, ensure that the IGV is focused on the correct loci coordinates within the correct genome, and also ensure that reads have not been downsized.

Step two for proper manual review of variants is to determine the total quantity of variant support. To quickly ascertain the total coverage, variant support, and variant allele frequency (VAF), you can either click on, or hover over, the loci of interest within the coverage track. This will create a popup window with information on the selected loci to provide a better understanding of the total amount of variant support within the selected track. This popup will detail if there are multiple variants (MV) (Figure S13), low count normal (LCN) (Figure S7), low count tumor (LCT) (Figure S8), low variant frequency (LVF) (Figure S10), or no count normal (NCN) (Figure S14). To modify popup text behavior in panels, you can click on the yellow comment icon in the Genome Ruler track to toggle between click and hover options. If available, you can also visualize this information for other tracks including relapse libraries, metastatic libraries, and RNA-sequencing libraries. Increased support in multiple tracks increases confidence in a true somatic call.

The third step is to evaluate the quality of the variant support in all tracks. First we directly visualize read strands that show variant support to determine the overall quality of the variant in question. This includes looking for read strands that have multiple mismatches (MM) (Figure S11) and high discrepancy regions (HDR) (Figure S6). We also look for read strands that are translucent or transparent, indicating low mapping quality (LM) (Figure S9). If variant support comes from low quality read strands, it increases our skepticism on the validity of the variant being evaluated. To quantify the mapping quality for read strands, you can click on individual read strands that show variant support. This will create a pop up window with information on read strand quality. You can also evaluate the base quality for individual bases showing variant support. On visual inspection, the base quality of the variant for a given read is reflected by the transparency of the letter. The majority of variant-supporting reads should have a high base quality and should not be translucent. Each infraction in variant support quality decreases our confidence that the variant in question is a true somatic variant. The final part of step three is to evaluate the normal track for tumor support in normal tracks (TN) (Figure S17) and normal track quality. Especially for hematologic tumors, it is important to evaluate the average level of tumor contamination across the normal samples to inform manual review decisions.

The fourth step requires checking for other sequencing artifacts. This requires first toggling between view as pairs (right click for each data track > click “view as pairs”) to visualise short inserts (SI/SIO) (Figure S15). You also must use the zoom in (+) and zoom out (-) buttons on the Genome Ruler track to visualize adjacent indels (AI) (Figure S1), high discrepancy regions (HDR) (Figure S6), multiple mismatches (MM) (Figure S11), same start end (SSE) (Figure S16), and ends (E) (Figure S5). Finally, it is recommended to evaluate the reference sequence in the Genome Features track to check for low complexity regions such as mononucleotide runs (MN) (Figure S12), dinucleotide runs (DN) (Figure S4), and tandem repeats (TR) (Figure S18). Evaluating the reference sequence is especially when the variant in question is a short insertion or a short deletion.

The fifth step requires synthesis of all of the available information to make a definitive call on the variant in question. To do this, we select a button on the IGVNav interface under the “Call” section. Calls are mutually exclusive and only one call can be associated with each variant. Step six requires selecting “Tags” to annotate each variant using the IGVNav interface. Any variant, even variants called as “somatic”, can be associated with a tag and multiple tags can be used to describe a single variant.

Finally, if the manual reviewer requires variant annotation that cannot be covered by the existing IGVNav features, we have provided a “Notes” section. This can be used to indicate large adjacent structural variants, dinucleotides, fusions, etc.

## RESULTS

### Understanding common sequencing patterns observed during manual review

Retrospective analysis of variants that have undergone somatic variant refinement through manual review allowed us to find common patterns that influence reviewer decisions. Selected examples were identified for each pattern and visualized using IGV screenshots (Figure 4-5, Figure S1-S18). Screenshots were annotated to emphasize aspects used to determine the eventual variant call or variant tag. Individuals proficient in manual review provided additional helpful hints to supplement screenshots based on their own findings and procedures. This includes warnings for challenging tumor types, IGV features used to elucidate sequencing patterns, and instances where there might be deviations from standard protocol. For example, when conducting somatic refinement on hematologic tumors, the allowable threshold for Tumor in Normal (TN) might be increased due to contamination of tumor cells within normal skin biopsies (Figure S17).

**Figure 4.**
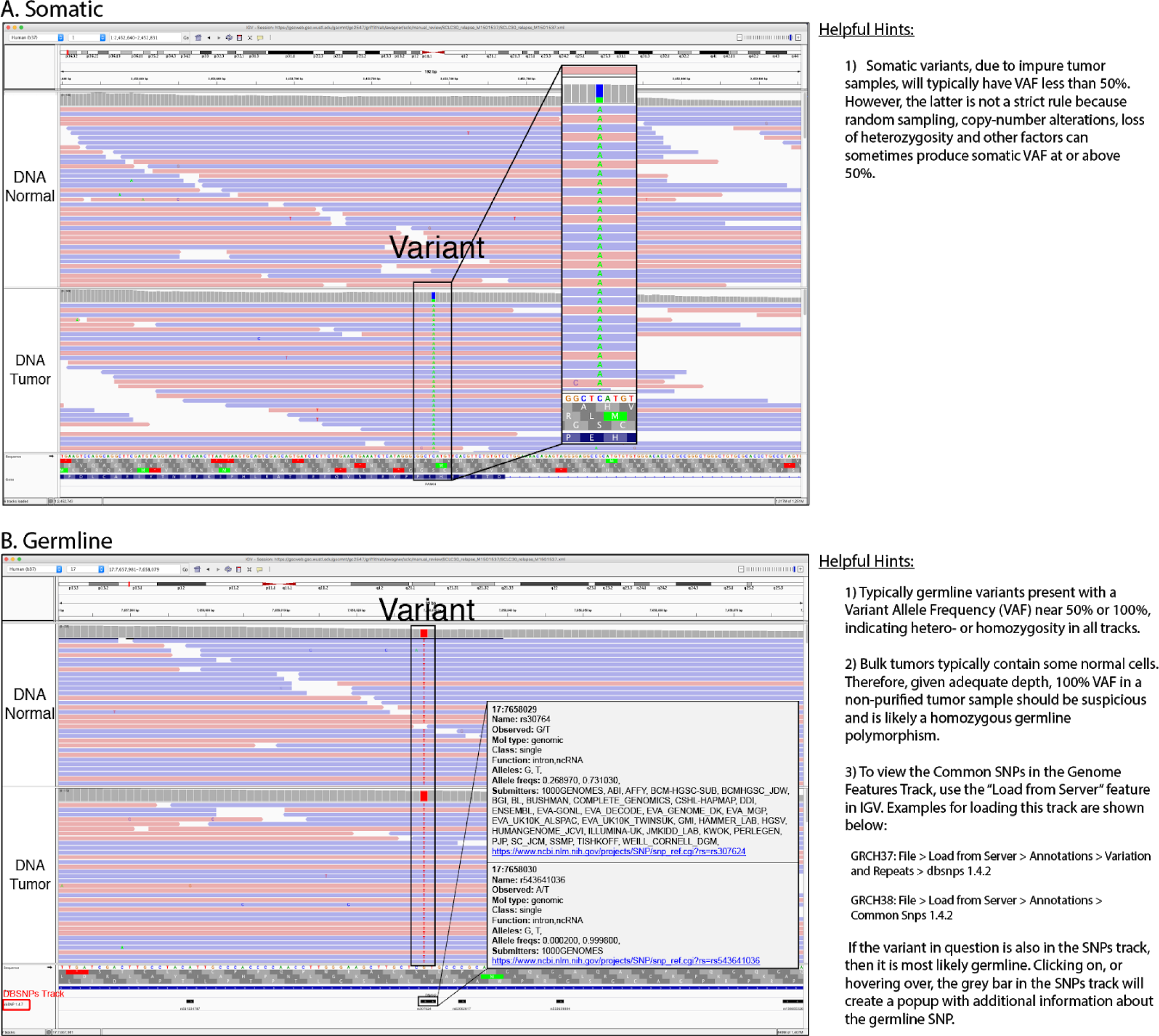
Example of calls that would be labeled as Somatic or Germline. **A.** In this example, the variant is presumed to be a real somatic variant. When evaluating the reference sequence in the Genome Features track, the reference allele is a cysteine (C). The alignments and coverage in the DNA Tumor track show that approximately 50% of reads that support a variant adenine (A) allele (green). Importantly, there are no reads supporting the variant in the normal sample, indicating that it is a somatic variant as opposed to a germline polymorphism. Using the gene annotation track, we can predict that this (C->A) base change would result in a TGG (W; Tryptophan) to TTG (L; Leucine) missense mutation in the RESP18 gene (note this gene is transcribed on the negative strand). **B.** In this example, the variants is presumed to be a germline polymorphism. The reference allele is an adenine (“A”), however in the DNA Normal track and the DNA Tumor track, all supporting reads are guanines (“G”). This indicates that the variant is likely a homozygous germline polymorphism. The Common SNPs track, provides further support that this loci is a common polymorphism.

**Figure 5.**
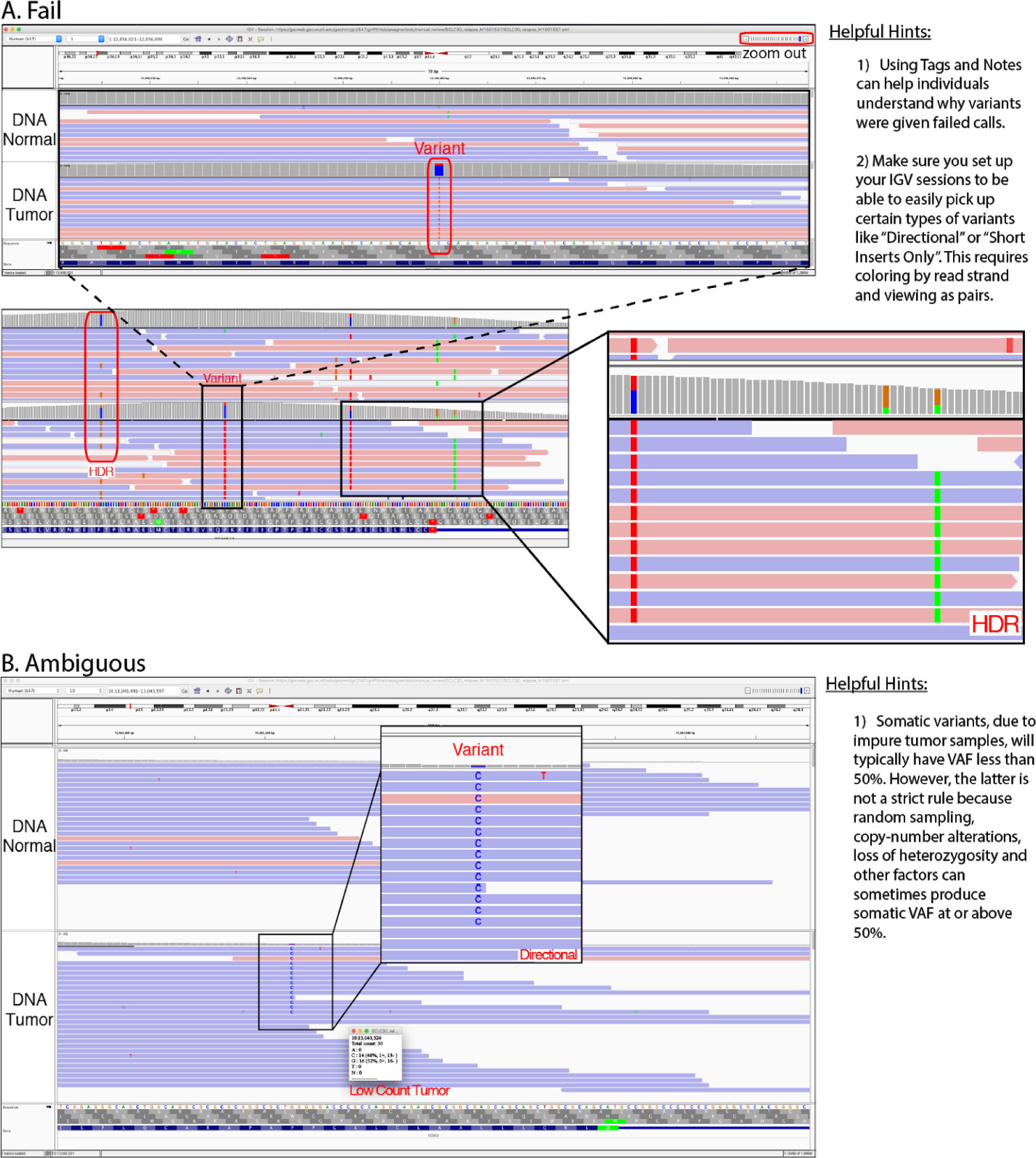
Examples of calls that would be labeled as Fail or Ambiguous. **A.** Ambiguous calls are made when the variant could be a real somatic variant, but the reviewer is not confident due to features of the variant called and corresponding reads. In this example, the variant has support from fourteen reads, but most are on negative read strands (93%). Additionally, several of the supporting reads have multiple mismatches indicating potentially low quality reads. However, there is a variant allele frequency (VAF) of 48%, which indicates that if the variant is a true somatic variant, then potentially 96% of tumor cells contain this variant. More information would be required to call this variant somatic or fail, therefore, the correct label is ambiguous. **B.** Failed calls are made when the called variant is most likely a false positive. In the first panel, IGV’s default visual makes the variant appear to be somatic, however, in the zoomed-out panel, we reveal a region of high discrepancy. High discrepancy regions (HDR) typically occur via improper alignment in regions of high homology across the genome or due to errors in the reference assembly. Given the HDR pattern observed, this variant is most likely a false positive and should be called as fail during manual review.

### Criteria used by manual reviewers to identify artifacts

To elucidate effective methods for proper manual review, we queried expert reviewers on features they considered important in annotating variants. When seven reviewers were asked to identify the top fifteen features they considered most important, 100% ranked tumor variant base quality as relevant, and 6/7 reported tumor VAF, tumor depth, tumor variant average mapping quality, tumor variant count, normal depth, and tumor variant direction bias (number of supporting reads on negative strand versus number on positive strand) to be important. There were 28 other features that were considered important for manual review by at least one expert reviewer (Table S1). All features that were identified by manual reviewers were taken into consideration integrated into the the standard operating procedure.

### Analysis of four types of calls

IGVNav was designed to support 4 variant calls: Somatic “S”, Germline “G”, Ambiguous “A”, and Fail “F”. For a call to be labeled as somatic, the variant must be an alteration in the DNA that is specific to the tumor, and not present within the germline (Figure 4A). Conversely, a germline variant is an alteration in the DNA, relative to the reference genome, that is present in both the tumor tissue and the germline tissue (normal track). Given a germline variant, we expect the variant allele frequency (VAF) to be near 100% or 50% in both the normal and tumor tracks, indicating homozygosity or heterozygosity, respectively. Sequencing depth should be kept in mind as sampling error impacts the accuracy of the VAF. It should be noted that labeling a variant as germline during manual review after being called somatic by a variant caller is suspect and might reveal underlying issues with the MPS pipeline being used (Figure 4B). To fail a variant, the reviewer must unequivocally determine that the variant was called because of a sequencing or alignment artifact. In the provided example, the called variant appears somatic in IGV’s default settings, however, when the variant is visualized in the context of a larger genomic area, a high discrepancy region is revealed to decrease confidence that the variant is a true positive (Figure 5B). If a reviewer has any residual doubt about failing a variant, then the reviewer should label the variant as ambiguous. This typically means that the reviewer would require more sequencing depth and/or better quality ready to make a confident call. An example of an ambiguous call is shown in Figure 5A, whereby there is no support for the variant in the normal track, and there are 14 reads supporting the variant in the tumor track. However, most of the reads are supported by negative read strands, some reads have multiple mismatches, and many of the supporting reads are short inserts (not visualized). While these examples provided insight into how variants are called, it is equally important to discuss the tags that can annotate the reasons behind variant calls.

### Analysis of nineteen types of variant tags

For each call made during manual review, especially for failed and ambiguous calls, it is important to classify the reason for the call using one or more of the nineteen tags available on the IGVNav interface. As mentioned, even though calls must be mutually exclusive (i.e. each variant is classified by only one call), a variant can be associated with multiple tags. Each tag represents a unique sequencing pattern or artifact that is commonly observed during manual review. These patterns can arise during DNA fragmentation, library construction, sequencing, or read alignment. Alternatively, some observable concerns can be caused by structural or complex issues intrinsic to the tumor being evaluated. Below, we describe how these concerning read strands are created within the sequencing pipeline and detail the resulting pattern observed in IGV.

The first step in the MPS pipeline is extracting nucleic acids from a tissue sample. The tumor type and tissue origin can play a role in generating artifacts that are observed when manually reviewing individual variants. For example, hematologic tumors (e.g. acute myeloid leukemia with high blast counts) can cause Tumor in Normal (TN) patterns due to infiltration of tumor cells in the normal biopsy. This can cause heavy variant support in the normal tracks and might cause confusion about the variant call (Figure S17). Generally, it is important to understand the average level of contamination across all variants within an individual tumor to determine an acceptable threshold for TN. This may require initial manual review of multiple variants prior to making definitive calls on any variants. Artifacts can also be generated during the tumor sample preparation. For example, Short Inserts (SI) or Short Inserts Only (SIO) are frequently observed when sequencing degraded nucleic acids (e.g. analysis of formalin fixed paraffin embedded (FFPE) samples).^27^ When generating paired-end reads, degraded and/or short molecules will produce two sequences that have overlapping alignments. This causes increased support for a variant, when in reality, it is merely the same molecule being evaluated multiple times (Figure S15). Short inserts can be visualized by viewing reads as pairs and looking for grey bands in the middle of read strands. Finally, given that many real variants are present at a low VAF, due to subclonal mutations or low purity tumors, Low Count Tumor (LCT) (Figure S8) and Low Variant Frequency (LVF) (Figure S10) can prevent a variant from confidently being called somatic. For example, if a tumor variant is only present in 10% of cells sequenced, the resulting VAF would be only 5%, such that reduced coverage in the tumor track (i.e. LCT) might preclude adequate variant coverage even though the variant might be somatic.^28^

Nucleic acid extractions are subsequently subjected to fragmentation, library construction, and optionally, hybridization capture. Additional errors can arise during this process. For example, a selection bias might skew which molecules are sequenced resulting in a lack of even distribution of sequencing across the desired genome space.^29^ These errors are labeled as: Low Count Normal (LCN), No Count Normal (NCN), and Low Count Tumor (LCT). No Count Normal (Figure S14) and Low Count Normal (Figure S7) are defined by no or few read strands in the normal tracks. Low Count Tumor (Figure S8) is defined by few read strands in the tumor track. Although the threshold for adequate coverage depends on the sample type and internal lab requirements, our lab require at least 20X coverage in both the tumor and normal tracks for a variant to be considered somatic. LCN, NCN, and LCT can occur if fragments that would map to the region are not adequately amplified during library construction and/or are not adequately sequenced. DNA quality and quantity, capture reagent balance and efficiency, and sample balance in multiplexed preparations can all impact the uniformity of coverage for a given sample.

After fragmentation and library preparation, the fragments are amplified using polymerase chain reaction (PCR). During PCR amplification, it is possible to introduce Directional (D) and Same Start/End (SSE) artifacts. Directional artifacts occur when variant support is only apparent on reads in a specific direction (i.e. either positive or negative strand). Typically, this occurs because the sequencing context affects the polymerase in one direction more than the reverse complement. Of note, directional artifacts can also be caused during alignment or during post-sequencing base/read processing (Figure S3). SSE artifacts occur when a small molecule is preferentially amplified and not removed through read deduplication programs.^30^ This artifact can be confirmed when all variant support reads have the same start and end position after alignment (Figure S16).

The next step in the MPS pipeline is sequencing. Sequencing errors are defined as nucleotides misread by the sequencing instrument, which can be caused by inefficiencies in sequencing chemistry, technical errors made by the camera system, and interference from neighboring clusters. One type of sequencing error is called “dephasing errors”, which occur when a nucleotide without a proper 3’ -OH blocking group is incorporated or the 3’ -OH group is not properly cleaved. This creates a situation whereby affected fragment(s) are out of sync with the cluster, contributing to background noise.^31^ One example of a dephasing artifact is labeled as Ends (E). These tags are used when variant support only occurs at the end of sequencing reads (within 10 base pairs), where there is an increased likelihood of dephasing error from chemical incorporation of blocking groups (Figure S5).

Finally, after reads are sequenced, they must be aligned to the reference genome. A large number of artifacts arise from poor alignment of sequence reads to the reference genome. These artifacts include: Mononucleotide Repeats (MN), Dinucleotide Repeats (DN), Tandem Repeats (TR), High Discrepancy Regions (HDR), Low Mapping (LM), Multiple Mismatches (MM), Adjacent Indels (AI), and Multiple Variants (MV). Many of these artifacts (MN, DN, TR) are attributable to regions of low complexity adjacent to the variant loci. Mononucleotide runs (Figure S12), Dinucleotide runs (Figure S4), and Tandem Repeats (Figure S18) typically occur when there is a base pair deletion or insertion adjacent to one, two, or three or more base pairs repeats, respectively. The remaining alignment artifacts (HDR, LM, MM, and MV) occur when single reads map to multiple or incorrect regions due to homologous sequences at multiple loci, highly variable regions between or within individuals (e.g., VDJ regions in immune cells), and errors in the reference genome. High Discrepancy Regions are apparent when multiple reads contain the same mismatches with the reference genome at various locations along the read strand (Figure 5B, Figure S6). Low Mapping can be determined by visualizing low mapping quality of reads containing the variant. Reads with mapping quality 0 (maps to multiple regions) are translucent when the reads are colored by read strand (Figure S9). MM is a label for variants that are supported only by reads that disagree with the reference genome at multiple loci across the same read indicating that the reads are of low quality or misalignment (Figure S11). Similarly, Multiple Variants, which is defined by read support for three or more bases at a given loci, indicates poor quality or misaligned read strands (Figure S13). Additionally, complex variants can induce false positives and create artifacts that are observed in IGV. Specifically, Adjacent Indels (AI) are observed when a large structural variant or small indels in a repetitive region causes local misalignment and creation of an apparent SNV or indel (Figure S1). All of these artifacts require careful scrutiny of the reference genome, base quality, and mapping quality.

In rare instances, we observed artifacts that were unable to be categorized and were labeled as Ambiguous Other (AO). In the example provided in Figure S2, the insertion variant showed high support in the tumor track, however, there were multiple variants at the loci being evaluated, multiple mismatches in reads showing variant support, and there was some tumor contamination in the normal track. Additionally, the variant might have been attributed to alignment issues since there was a tandem repeat sequence adjacent to the varant. Finally, many reads in both the normal and tumor tracks had strange ends in non-variant supporting reads indicating potential sequencing concerns at the genomic loci. Given that this sequencing pattern is non-descriptive, it is recommended to include a short note justifying the reason for the Tag and eventual variant Call.

## DISCUSSION

We hope that the use of the presented manual review standard operating procedure will improve the refinement of putative somatic variants after somatic variant calling. Our manual review protocol creates a method to accurately filter false positives and call true positives from a list of variants identified as true from automated somatic variant callers. Our outline provides a method to properly label variants based on common sequencing artifacts observed during the manual review process. It also provides methods to ensure that all artifacts are observed by creating a standard method for IGV setup and navigation through the variants of interest.

Use of the IGVNav software allows for standardization of the input and output files for somatic variant refinement during manual review. Its features, including Calls, Tags, and Notes, create distinct labels for each variant and allow for retrospective analysis of sequencing quality across tumor cohorts. The output file can also be used as an input file for databases that can annotate variants based on clinical significance. For example, the output file from IGVNav can be filtered for true positives and compared to the CIViC database^32^ to quickly assess variants that have clinical significance. These variants can be mapped to the CIViC Application Programming Interface (API) to determine sensitive and resistant therapeutics as well as diagnostic, prognostic, and predisposing implications.

It is our intent to continuously improve this manual review protocol through subsequent revisions. As we continue to use the manual review process for elimination of false positives, we learn new ways to accurately elucidate somatic variants. We will also improve the protocols through incorporation of new tools that assist in the somatic variant refinement process. This will become increasingly important as we begin developing new ways for somatic variant refinement such as the adoption of machine learning algorithms to classify somatic variants. The use of machine learning algorithms could dramatically reduce the need for manual review, whereby only variants classified as ambiguous would require direct visualization.

There are intrinsic limitations associated with the manual review process that will not be rectified by use of this standard operating procedure. First, manual reviewers have reported fatigue after many hours of manual review. This is especially important when reviewing tumors with high variant burden such as melanomas or lung cancers. It is recommended that individuals schedule hour sessions and breaks between sessions to improve the consistency of manual review for the entire tumor or study cohort. Second, despite training individuals to review variants in a similar manner, there will also exist inter-reviewer variability. This variability will be especially true for variants determined to be ambiguous. Finally, manual review of variants might change over time as an individual begins to recognize the idiosyncrasies associated with particular tumor subtypes. For example, one might change the acceptable threshold of tumor contamination after reviewing many variants in an individual tumor. This is especially important for hematologic tumors or tumors with high level of tumor infiltrate. Finally, the scope of this SOP is limited to the manual review of somatic SNVs/Indels for somatic variant refinement, however, many of the aspects of the protocol, including setup and assessment, can be directly applied to other analyses, such as germline variant calling.

Many of the existing limitations of somatic variant refinement through manual review could be addressed through automation of post-processing of somatic variants. This would further standardize and systematize somatic variant refinement and reduce the labor burden required to develop a list of putative somatic variants for downstream analysis. Advancements in computing and improved cooperation and openness presents an opportunity for the existence of such a process.

## Author Contributions

EKB wrote the manuscript, the manual review standard operating procedure, and conducted/analyzed all experiments. KMC, KK, PR, BJA, CR, FG, SLS, and LT wrote the manuscript. MM and AW wrote and updated the IGVNavigator tool. LT, KK, ZLS and BJA developed the initial Medical Genomics Manual Review Guidelines. SJS, MG, and OLG supervised the project and revised the paper.

## Acknowledgements

EKB was supported by the National Cancer Institute (T32GM007200-42). BJA was supported by the Siteman Cancer Center. SJS is funded by the National Library of Medicine (NIH NLM R01LM012222 and NIH NLM R01LM012482). MG is funded by the National Human Genome Research Institute (NIH NHGRI R00HG007940). OLG is funded by the National Cancer Institute (NIH NCI K22CA188163 and NIH NCI U01CA209936).

## Supplementary Tables and Figures

**Table S1.** Features labeled as important when surveying seven manual reviewers. When given a list of 71 features, expert manual reviewers were asked to prioritize the top fifteen features considered most important. A description of the features, listed in order of importance, is included in the Feature column and the number of reviewers listing each feature as important is included in the Reviewer Count column.

| Feature | Reviewer Count |
| --- | --- |
| Tumor Variant Average Base Quality | 7 |
| Normal Depth | 6 |
| Tumor Variant Allele Frequency | 6 |
| Tumor Depth | 6 |
| Tumor Variant Average Mapping Quality | 6 |
| Tumor Variant Count | 6 |
| Tumor Variant Number Minus Strand | 6 |
| Tumor Variant Number Plus Strand | 6 |
| Normal Variant Allele Frequency | 5 |
| Normal Variant Count | 5 |
| Tumor Reference Count | 5 |
| Normal Reference Count | 4 |
| Tumor Other Base Count | 4 |
| Tumor Reference Average Base Quality | 3 |
| Tumor Reference Average Mapping Quality | 3 |
| Tumor Reference Number Minus Strand | 3 |
| Tumor Reference Number Plus Strand | 3 |
| Tumor Variant Average Number of Mismatches | 3 |
| Tumor Variant Average Singe End Mapping Quality | 3 |
| Normal Reference Average Base Quality | 2 |
| Normal Reference Average Mapping Quality | 2 |
| Normal Variant Average Base Quality | 2 |
| Normal Variant Average Mapping Quality | 2 |
| Tumor Reference Average Number of Mismatches | 2 |
| Tumor Variant Average Distance to 3' End | 2 |
| Normal Other Base Count | 1 |
| Normal Reference Average Distance to 3' End | 1 |
| Normal Reference Number Minus Strand | 1 |
| Normal Reference Number Plus Strand | 1 |
| Normal Variant Average Position as a Fraction | 1 |
| Normal Variant Average Singe End Mapping Quality | 1 |
| Reviewer | 1 |
| Tumor Reference Average Singe End Mapping Quality | 1 |
| Tumor Variant Average Clipped Length | 1 |
| Tumor Variant Average Position as a Fraction | 1 |
| Tumor Variant Average Sum of Mismatch Qualities | 1 |

**Figure S1.**
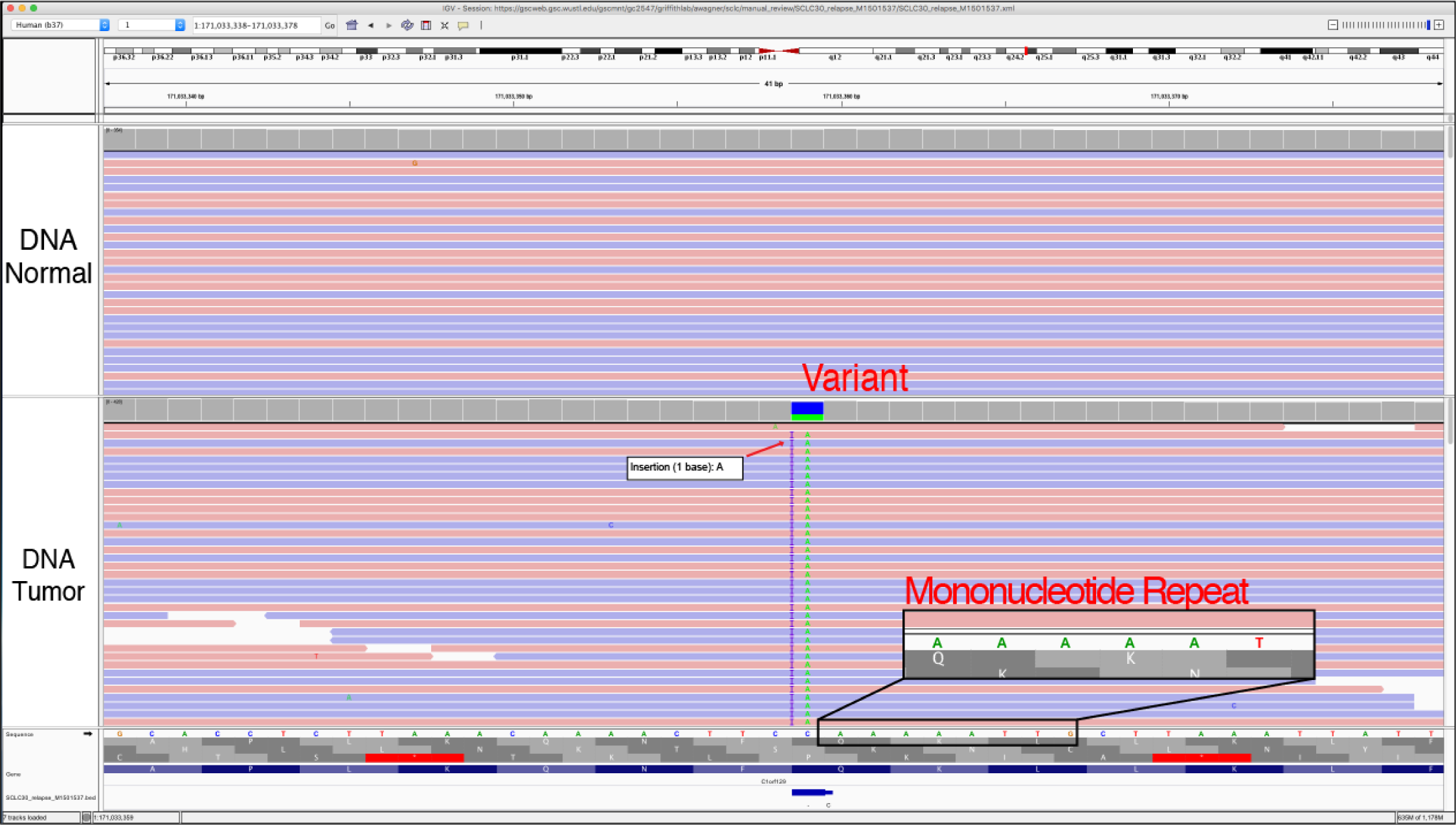
Example of an Adjacent Indel (AI). The Adjacent Indel Tag is used when a false positive somatic variant results from misaligned reads around a germline or somatic insertion or deletion (indel). In this case, the misalignment is occuring in the tumor DNA, due to the adjacent insertion, possibly due to a mononucleotide repeat near the adjacent indel and called somatic variant.

*Helpful Hints:*

1) To adequately catch this artifact, it is necessary to zoom out on the IGV session to ensure that you visualize the adjacent insertion or deletion.

2) It is important to evaluate the Genome Features track visualize possible tandem repeats that might be implicated in the misalignment.

**Figure S2.**
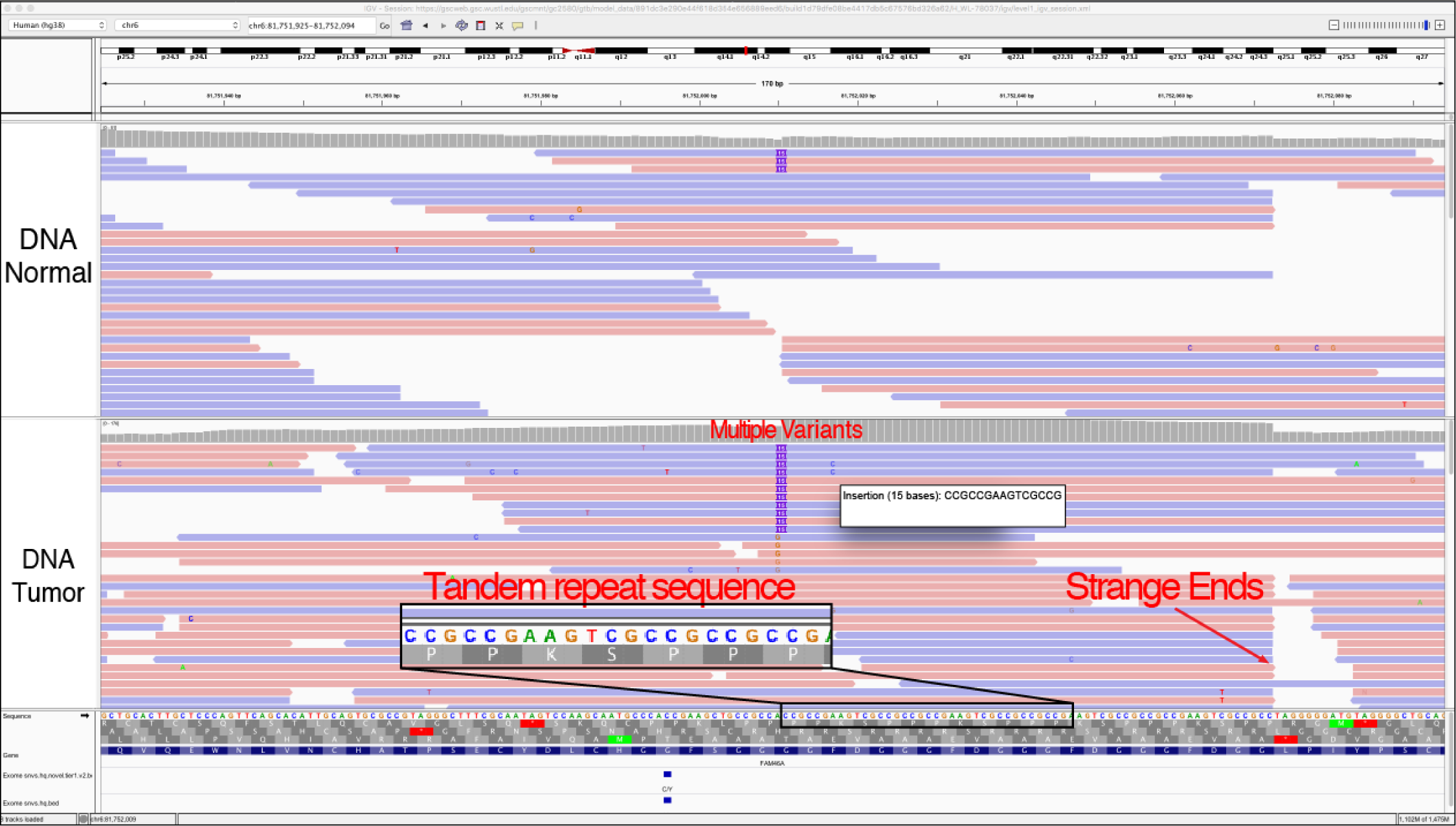
Example of an Ambiguous Other (AO). Ambiguous other is used to define a variant surrounded by inconclusive genomic features that can’t be explained by the other tags available. For example, genomic regions with increased A/T or G/C content that are not contained within tandem or dinucleotide repeats. This could also be described as a low complexity region. If the Ambiguous Other tag is used, it is highly recommended to include a short description in the Notes section.

**Figure S3.**
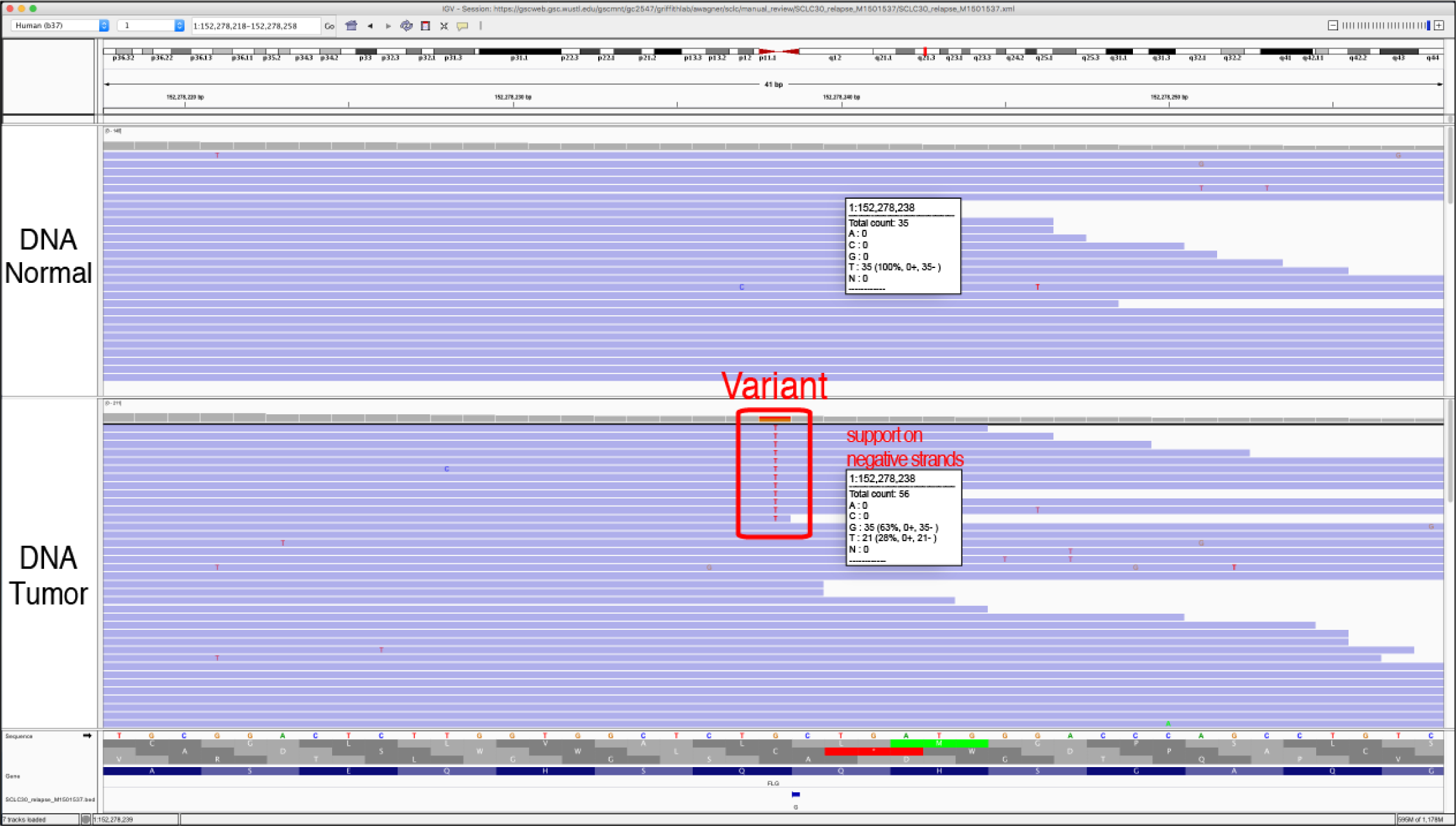
Example of a Directional (D). A directional artifact is when the variant being evaluated can only be found on reads that are sequenced in either the forward or the reverse direction. Typically, this is caused by strand bias during sequencing. To properly visualize the directional artifacts, you must make sure the IGV tracks are colored by read strand.

*Helpful Hints:*

1) This call can only be made when the reads are not viewed as pairs. When viewing data tracks as pairs, the reads in both directions are condensed and could possibly make the variant appear to be exclusively supported by read strands in a particular direction.

2) To adequately catch this artifact, it is necessary to color the alignments by read strand:

Right click on the track you want to color > click *“*Color alignments by*”* > click *“*read strand*”*

**Figure S4.**
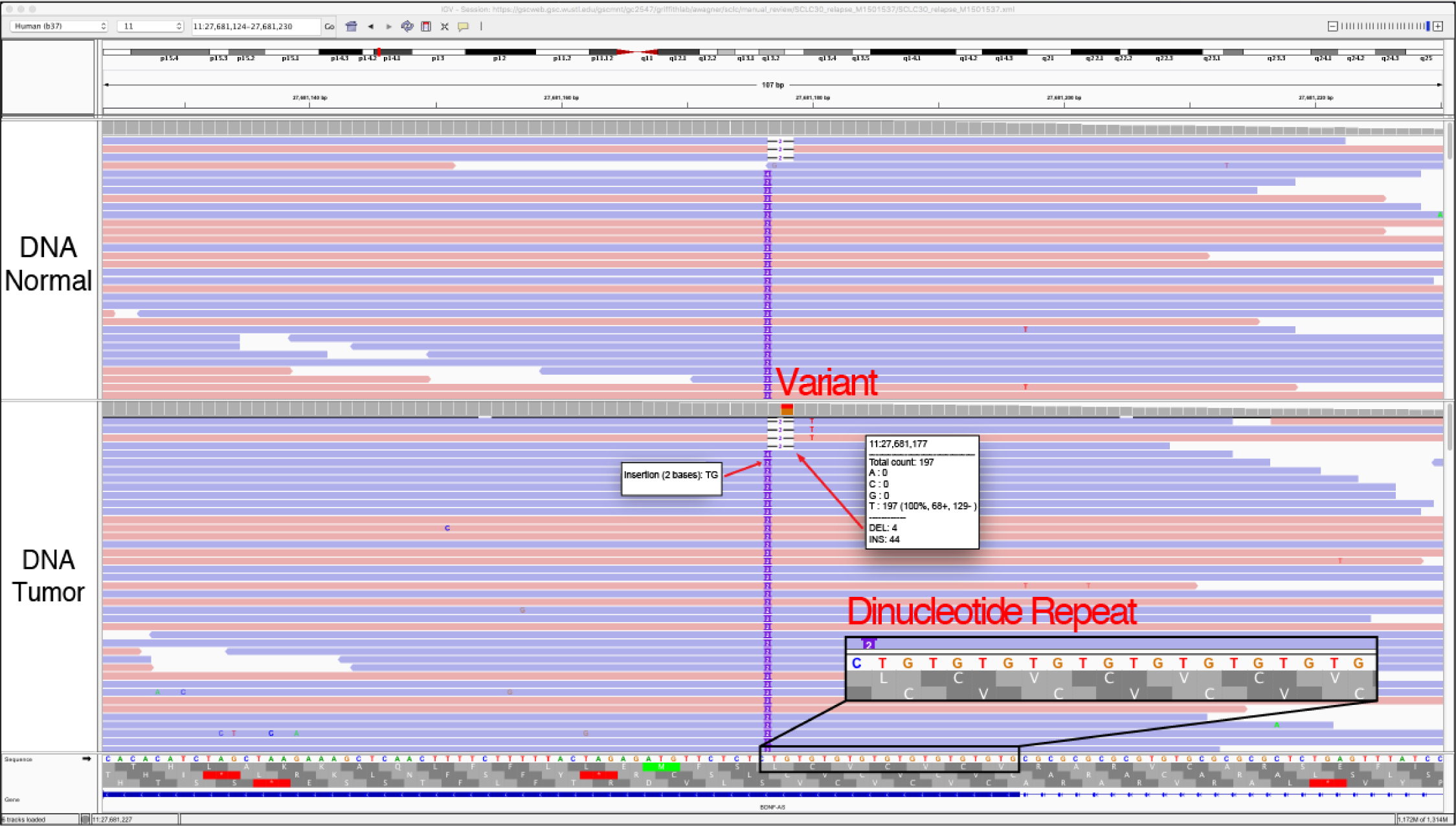
Example of a Dinucleotide Repeat (DN). The dinucleotide repeat pattern occurs when the called variant is adjacent to a region of the reference sequence that has two alternating nucleotides (e.g. TGTGTG…). In this instance, the called variant is most likely caused by misalignment of the read to the reference genome. Repeat regions are also areas where some sequencers, particularly those dependent on the polymerase enzyme, are prone to making mistakes. However, it is important to note that dinucleotide repeat are also a common source of normal human variation (inherited germline, de novo germline, or somatic) that arise due to errors produced by polymerase during DNA replication. Other factors, such as the size of the repeat, or appearance in the normal, should be considered during manual review to confidently call the variant. Special alignment, assembly, or even additional sequencing technologies may be needed to validate short repeats of this nature.

*Helpful Hints:*

1) Typically, these variants are small deletions or small insertions and they are usually visualized in the both the tumor tracks and the normal tracks.

2) Although the variant being evaluated may be a two base-pair deletion, deletions or insertions of different sizes are also often observed.

**Figure S5.**
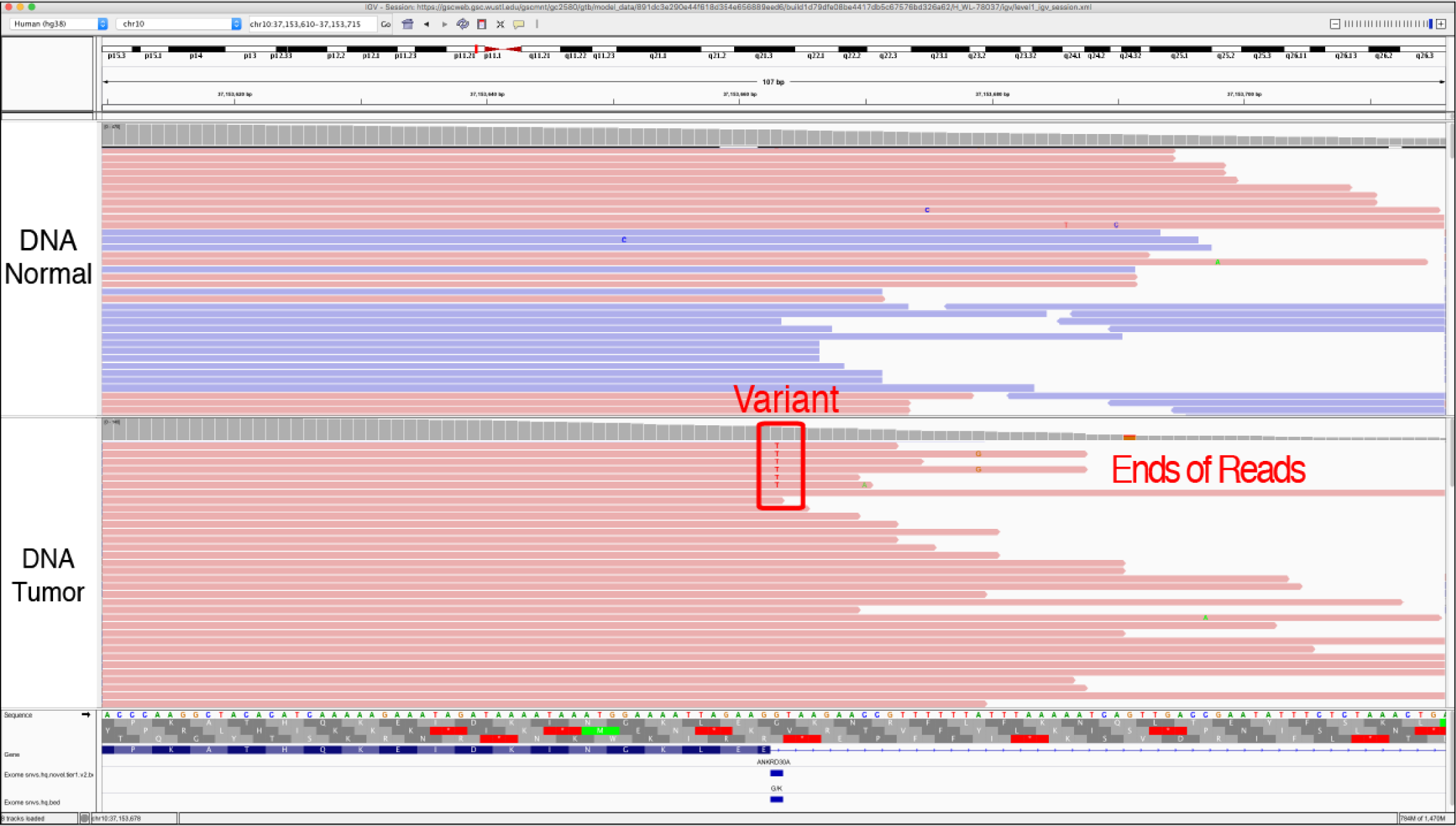
Example of an Ends (E). The variant called is only present close to the end (within 15 base pairs) of the variant-supporting reads. At read strand ends, there is an increase rate of error generation that can cause appearance of an erroneous variant. The multiple mismatches on the read strands that support the variant further support that this variant was due to a sequencing artifact.

*Helpful Hints:*

1) To adequately catch this artifact, it is necessary to zoom out on the IGV session to ensure that you visualize the ends of the reads.

2) This artifact must also be evaluated by coloring the alignments by read strand:

Right click on the track you want to color > click

*“*Color alignments by*”* > click *“*read strand*”*

**Figure S6.**
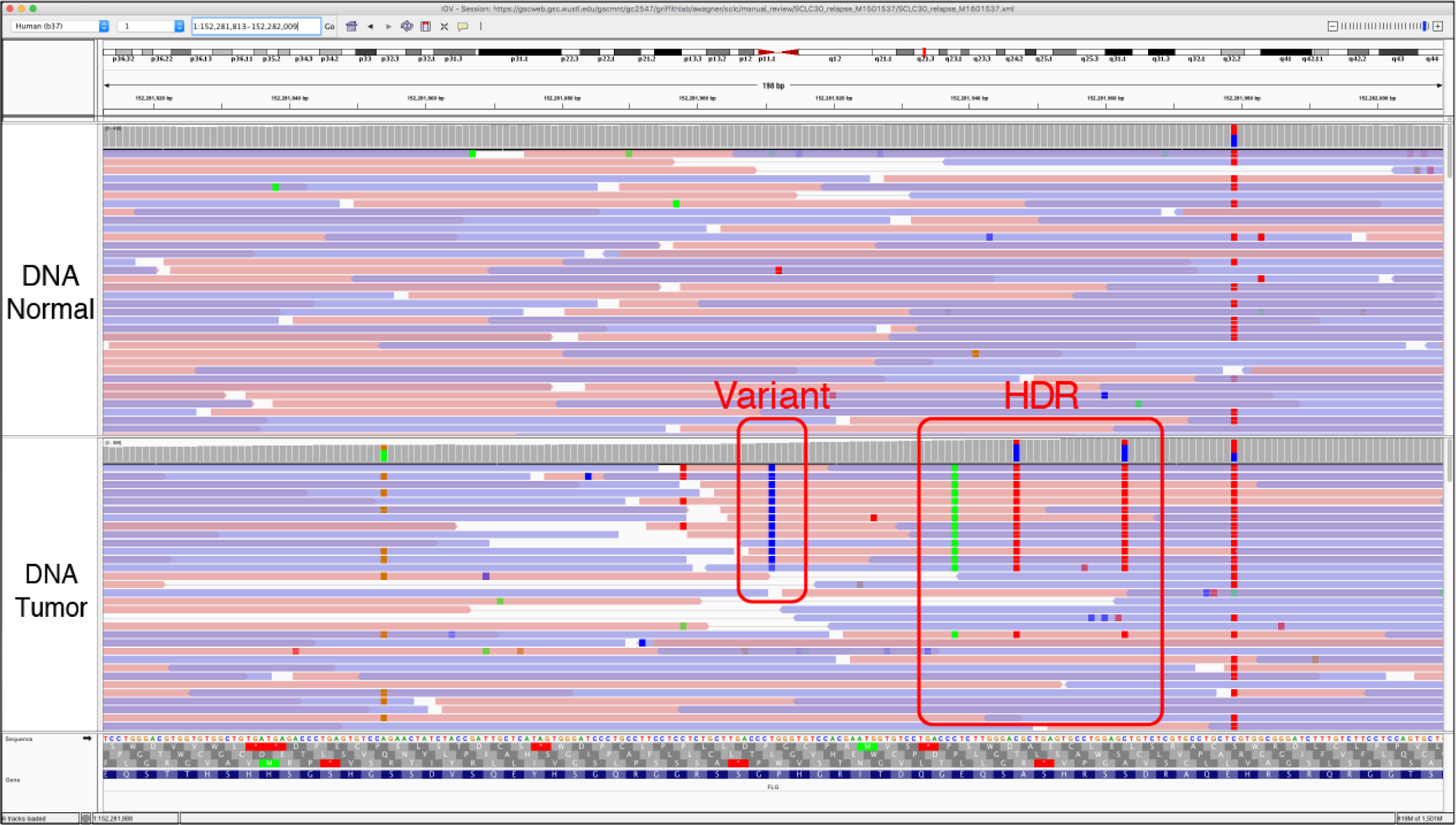
Example of High Discrepancy Region (HDR). The HDR tag is used when a variant is supported by reads that also have other mismatches that appear across the whole track or in multiple tracks. HDR typically occurs when there are homologs across the genome and mismapping of the reads causes an apparent variant when in reality, the variant represent differences between the homologs. The HLA locus and other highly polymorphic and duplicated loci are especially prone to this issue. These regions may require specialized alignment or assembly strategies for high quality variant calling.

*Helpful Hints:*

1) This tag is distinguished from Multiple Mismatches (MM) by the similarities of the mismatches across multiple tracks. In this example, all tracks contain the exact same mismatches at the same loci in the genome.

2) If there are multiple variants in a row that only 10-20 bases apart in the same gene then you should zoom out and make sure that you are not within a high discrepancy region.

3) It is important to be sure that the variant being evaluated is not due to a cluster of single nucleotide polymorphisms (SNPs). Sometimes, common SNPs can happen and be real and might be confused with an area of HDR. To distinguish the two it is important to consider the following: it is unlikely that a truly somatic variant would be observed on both alleles of a heterozygous SNP; therefore, reads supporting a variant should also support only 1 allele of the heterozygous SNP (be in linkage with one allele). This is another instance when having a track identifying common polymorphisms can be helpful.

**Figure S7.**
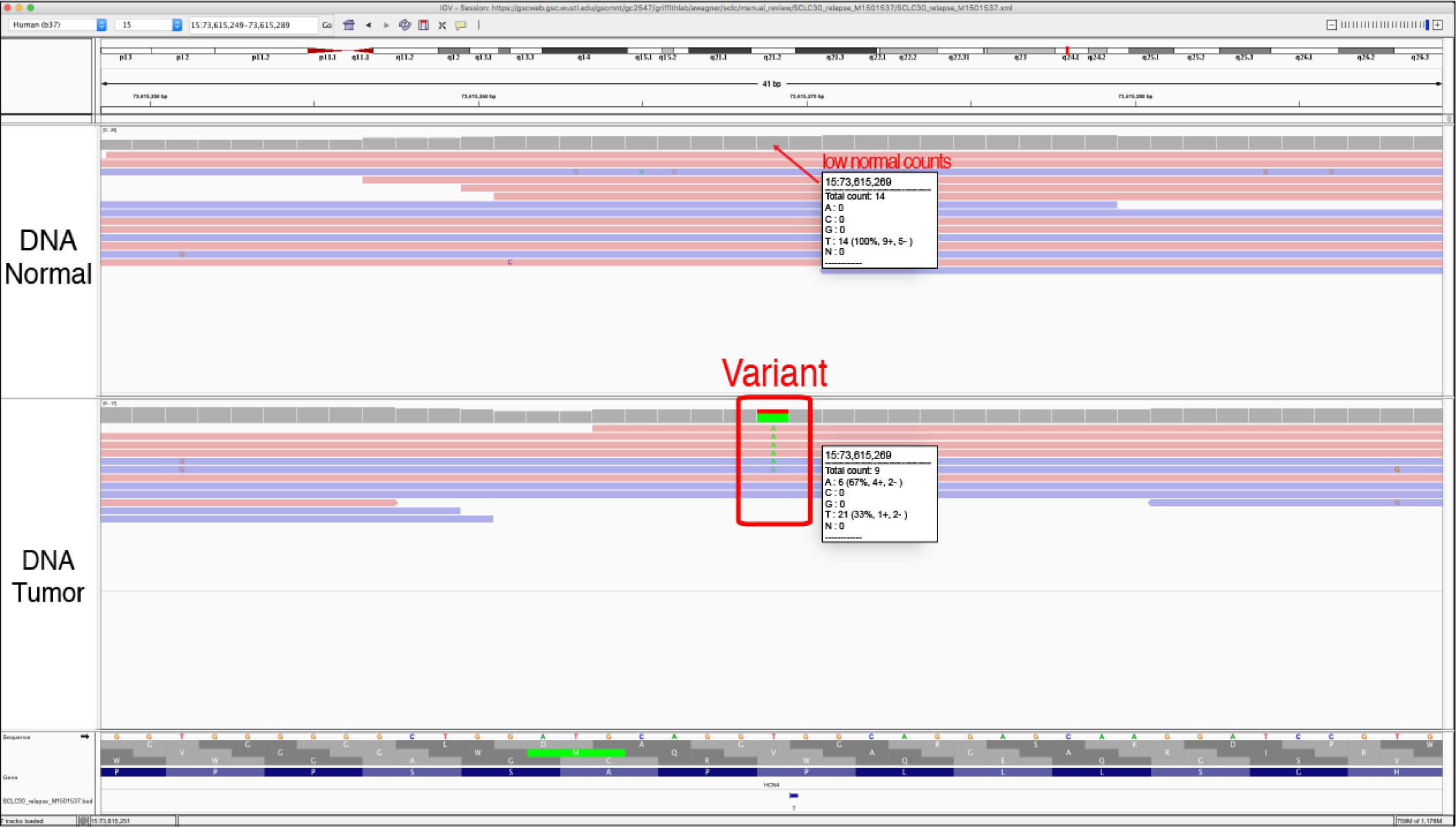
Example of Low Count Normal (LCN). The low count normal tag is used when there is not adequate coverage for the loci being evaluated in the normal track (coverage < 20X), which prevents effective comparison to the tumor sample. The insufficient coverage prevents us from making a confident somatic variant call. Coverage information can be assessed by clicking on the loci in the coverage track to reveal a popup with information about the reads at this loci. This threshold is experiment-specific, however, we typically require at least 20X coverage in both tumor and normal to be sure that a manual review call is a true somatic variant.

*Helpful Hints:*

1) If a variant has low coverage in the normal track, it can be treated like a “tumor only” sample. This might require populating the genomic features section with a common SNPs track to ensure that the variant is not a polymorphism.

**Figure S8.**
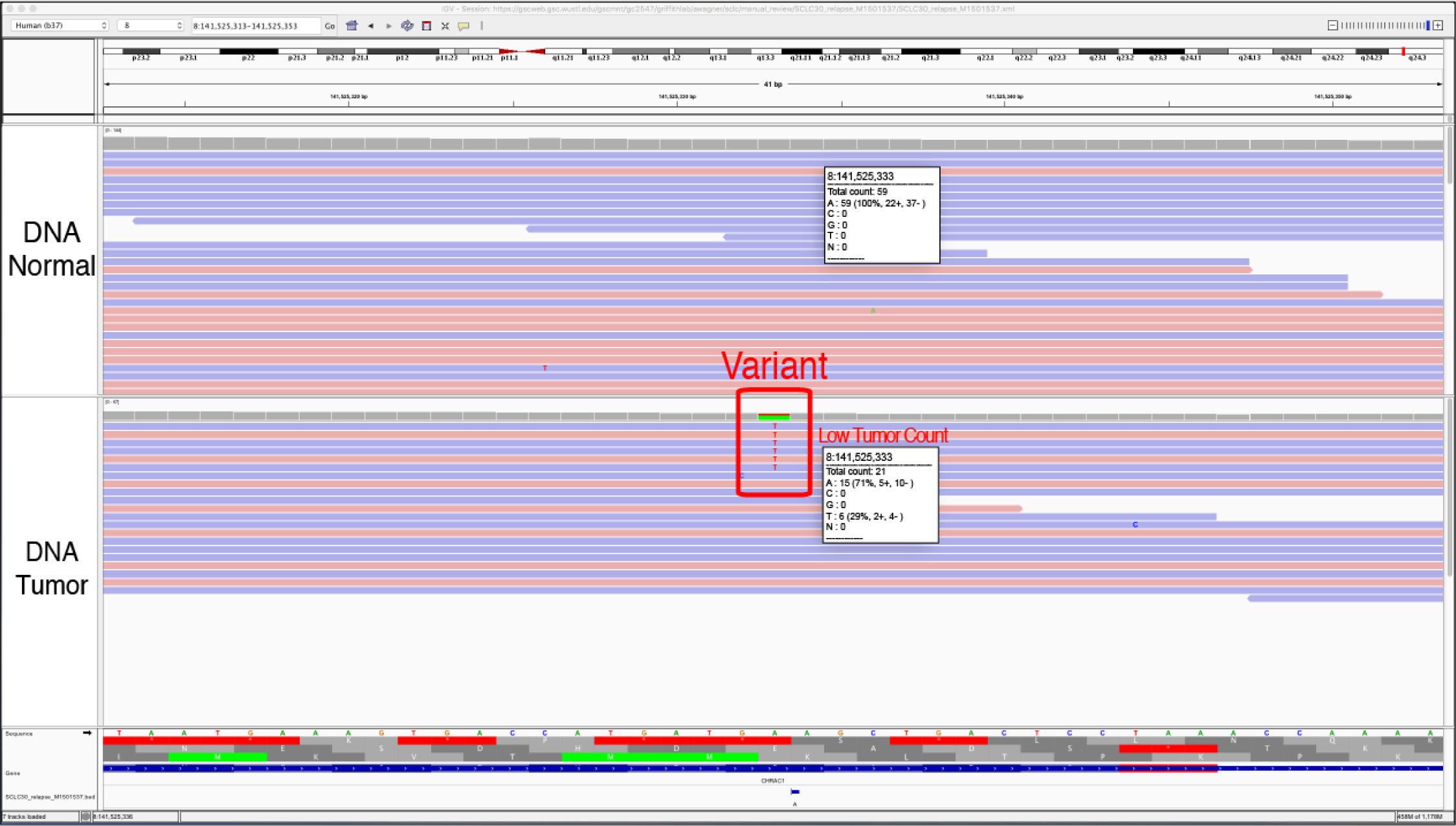
Example of Low Count Tumor (LCT). The Low Count Tumor tag is used when the coverage in the tumor is lower than the average. This threshold is experiment-specific. Low coverage in the tumor decreases the accuracy of variant allele frequency (VAF) estimates because sampling noise and uncertainty play a bigger role than if the coverage is very high. This may result in: 1) a false negative with an underestimated VAF, 2) a false positive due to over-estimation of the VAF, or 3) a true positive call with inaccurate VAFs.

*Helpful Hints:*

1) If a variant with a low VAF is determined to be somatic, it is important to consider this when employing downstream analysis. For example, the observed VAF might not be an accurate representation of the true VAF of the variant and therefore should not be used to make clinical judgements about the tumor.

2) Thresholds can be used pre-filter variants with low VAF in tumor or normal to eliminate the need to evaluate these variants during manual review.

**Figure S9.**
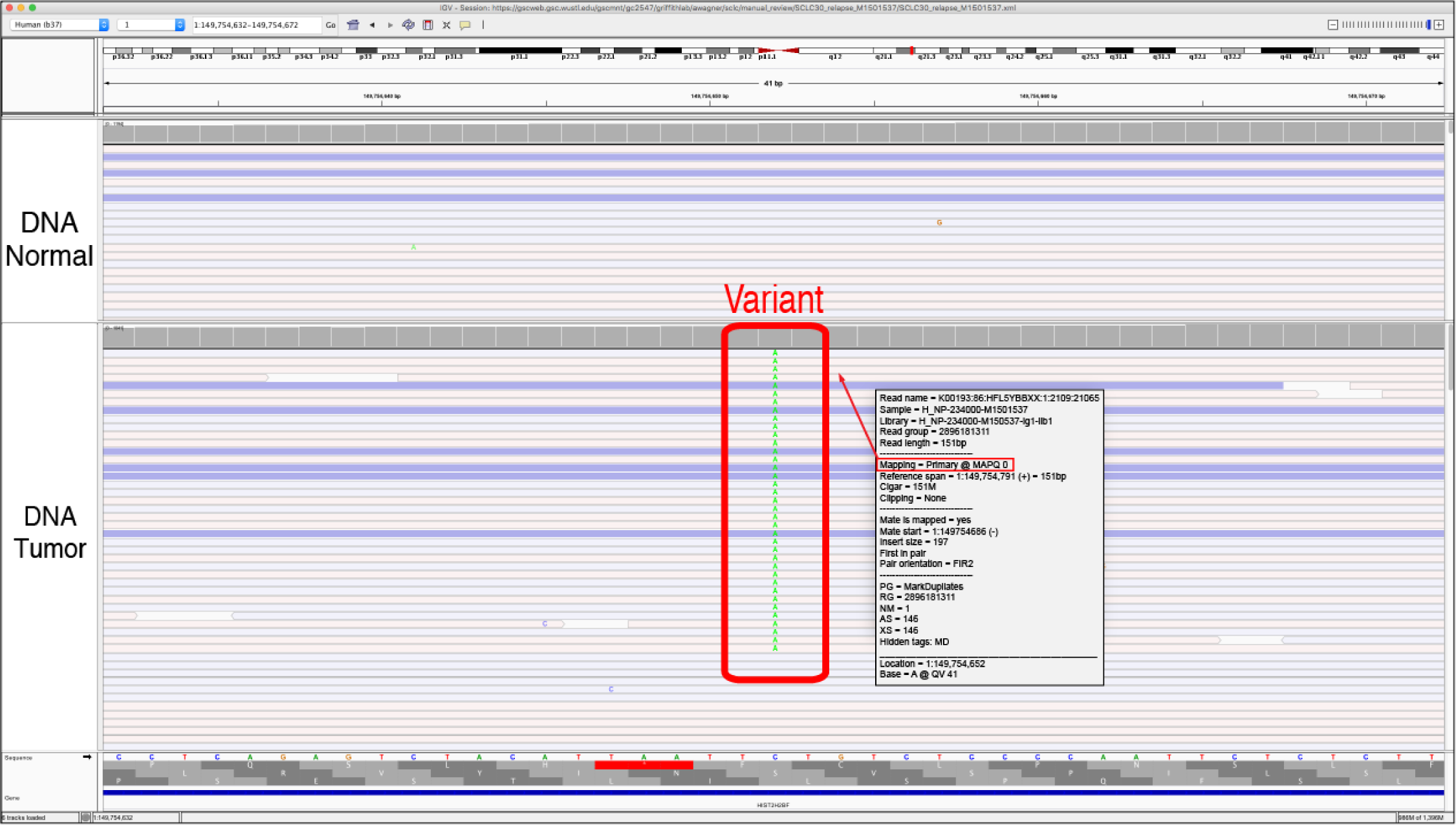
Example of Low Mapping (LM). Low Mapping is used to tag variants that are mostly supported by reads that have low mapping quality. The mapping quality of a read, when the reads are “colored by strand,” is indicated by the opacity of the read whereby lighter reads have lower mapping quality and darker reads have higher mapping quality. Mapping quality refers to a measure of confidence or probability that a read has been correctly aligned to the reference genome. This value can also be obtained by clicking on a read for its information. Variants that are supported primarily or solely by low mapping quality reads are considered suspect and tagged with LM.

*Helpful Hints:*

1) In regions where numerous reads have a mapping quality of 0, the reads are often mapped to multiple locations across the genome. This results in low mapping quality reads in both the normal and tumor. By right clicking on the read, you can visualize other mapping locations of individual reads.

2) By default, all reads are show in IGV, even if the mapping quality is 0. This threshold can be adjusted to eliminate low quality reads from IGV during manual review.

View > Preferences > Alignments > Mapping Quality Threshold

**Figure S10.**
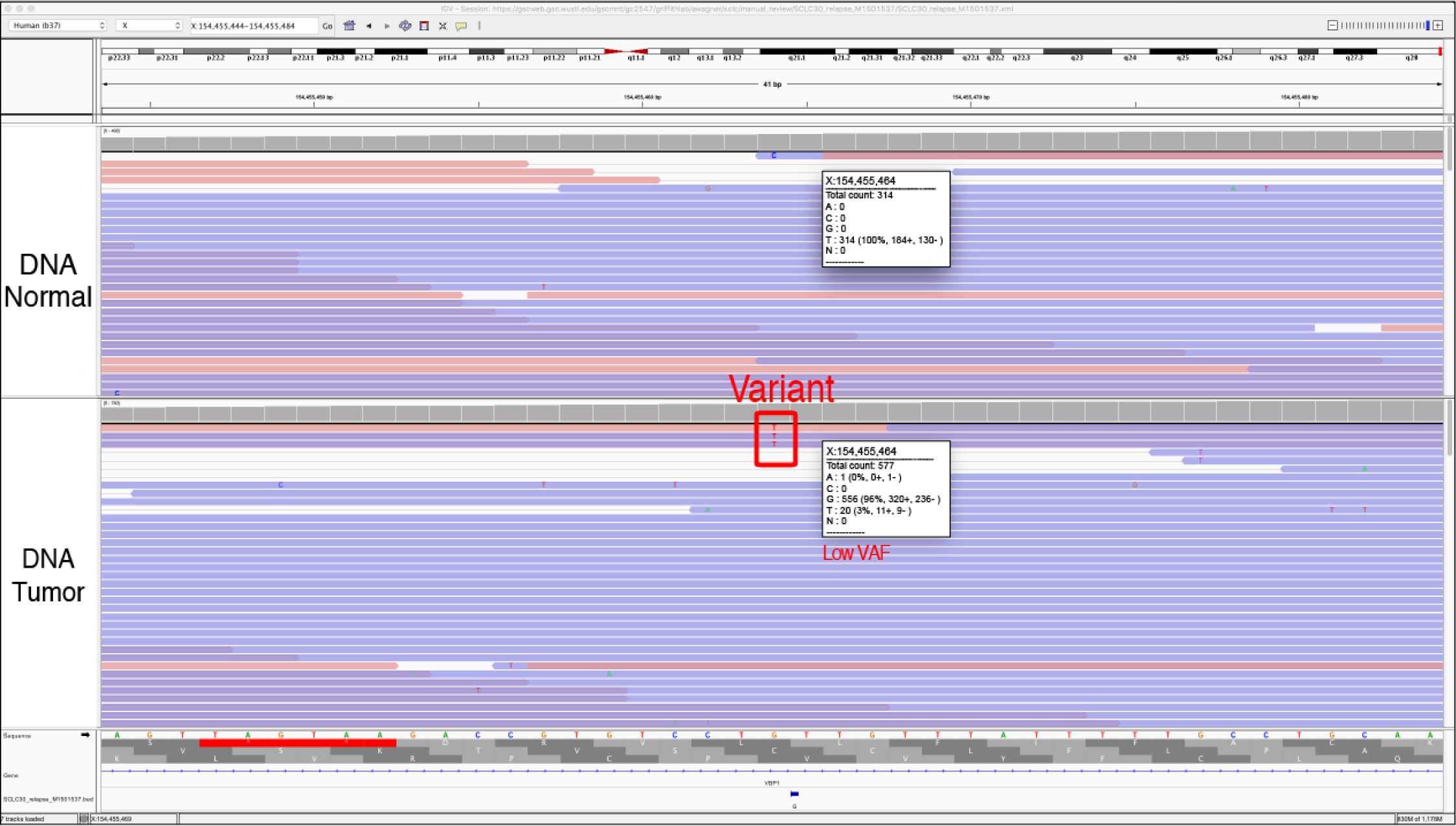
Example of Low Variant Frequency (LVF). The Low Variant Frequency tag is used when there are some reads of support for the variant, but the variant allele frequency (VAF) is relatively low. To quickly assess the VAF, you can click on the coverage track to pull up the total read counts, the number of reads for each base, and if the variant support is located on reads in the positive and negative direction.

*Helpful Hints:*

1) The coverage track will be colored according to base when a variant is present at a 15% VAF, by default. This threshold can be changed by using the following commands:

View > Preferences > Alignments > Coverage allele-fraction threshold

This can be particularly helpful with high depth samples and/or when low VAF (e.g., sub-clonal) variants are expected.

**Figure S11.**
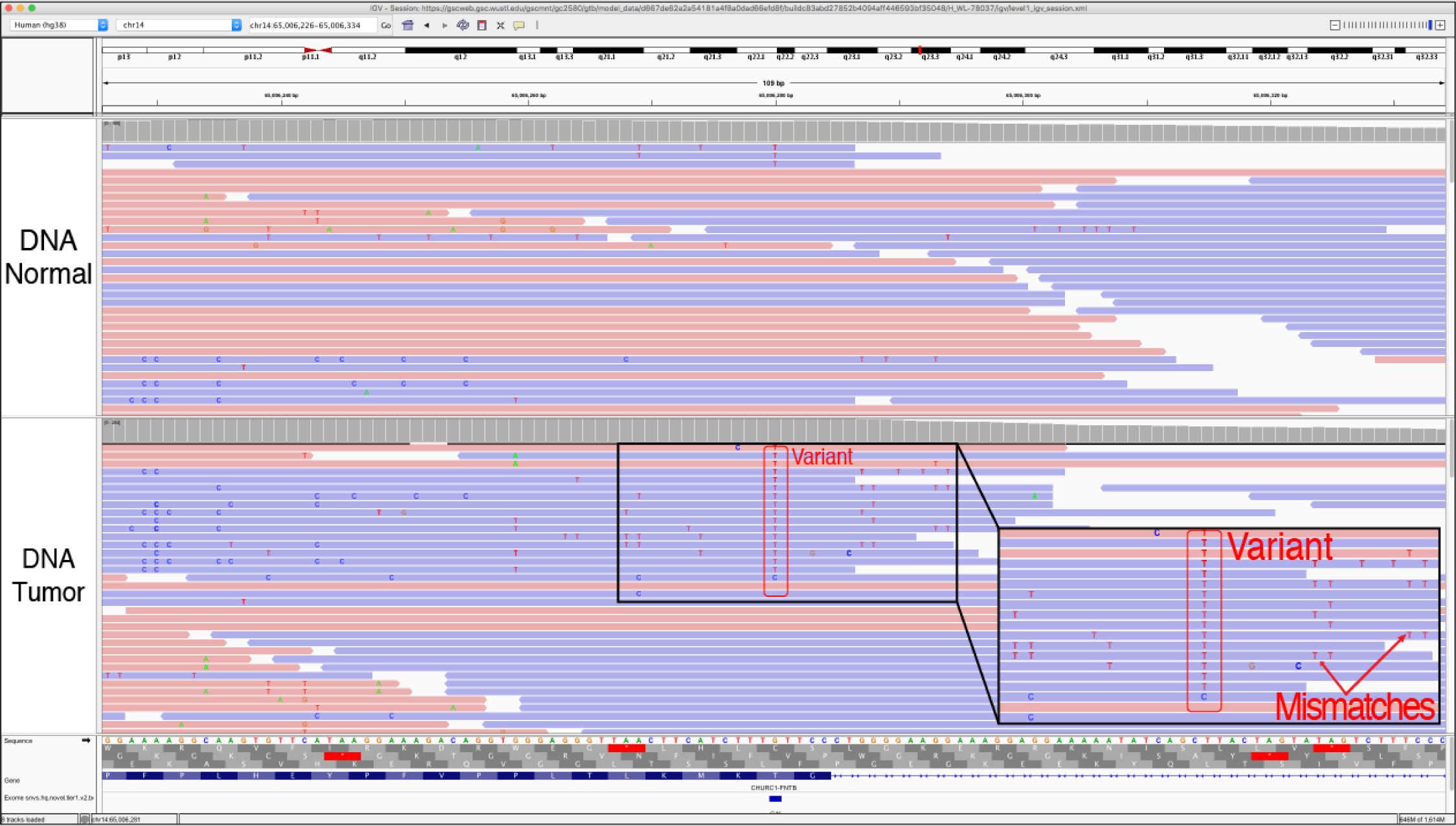
Example of Multiple Mismatches (MM). The multiple mismatches tag is used when the reads that contain the variant have other mismatched base pairs, which reduces the confidence in the read quality. This tag is similar to the HDR tag, however, it can include mismatches that are not exactly the same across multiple reads.

*Helpful Hints:*

1) The mismatch base color becomes more transparent as the base quality gets lower, so if the adjacent mismatch is darker in color, then you have reduced confidence in that read being properly sequenced and/or aligned to the reference genome.

**Figure S12.**
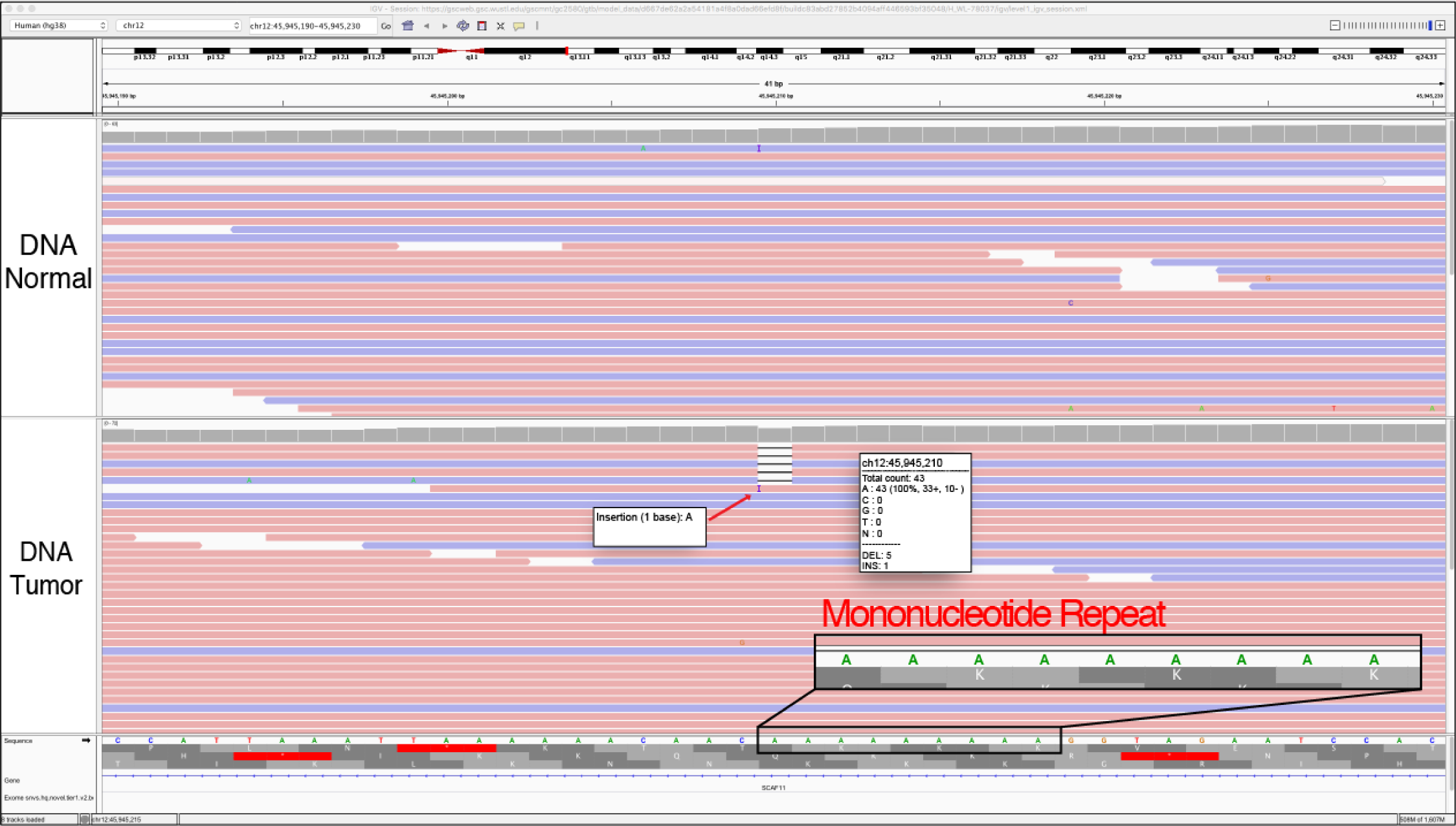
Example of Mononucleotide (MN). The mononucleotide repeat (homopolymer tract) tag is used when a variant is called in proximity to a region containing a a single nucleotide repeat (e.g. AAAAAAA…). The variant being evaluated may be a SNV or indel directly adjacent or within this region and called due to misalignment. Repeat regions are areas where some sequencers, particularly those dependent on the polymerase enzyme, are prone to making mistakes. However, it is important to note that these are also areas of normal human variation or real *de novo* mutations due to errors produced by polymerase during DNA replication. Like the Dinucleotide (DN) tag, other factors such as the size of the repeat, appearance in the normal and indels of varying length should be considered during evaluation.

*Helpful Hints:*

1) Typically, these variants are small deletions or small insertions and they are usually visualized in the both the tumor tracks and the normal tracks.

2) Although the variant being evaluated may be a 1bp deletion, deletions of different sizes or even insertions are often observed with artifacts.

**Figure S13.**
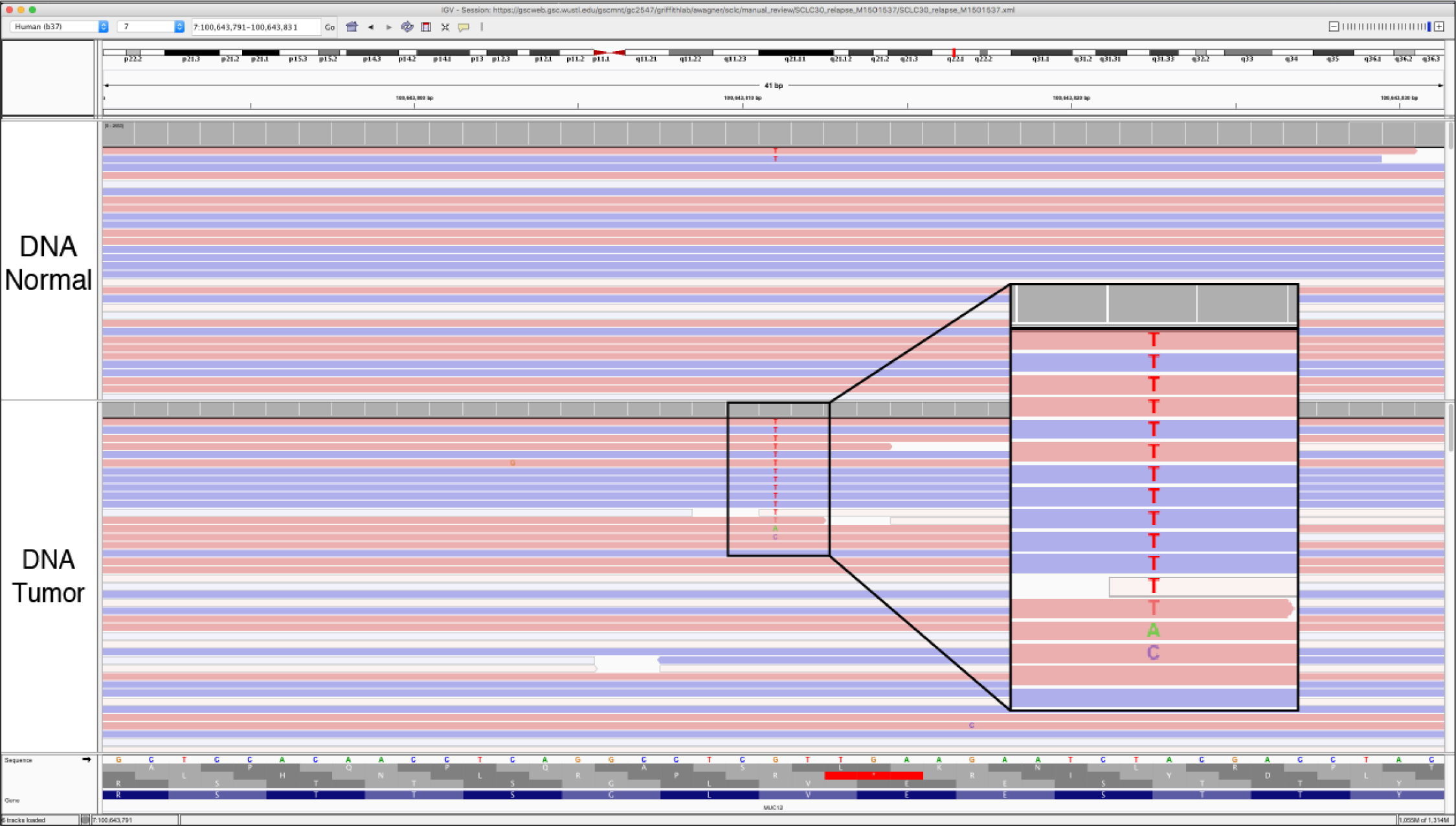
Example of Multiple Variants (MV). The Multiple Variants tag is used If the variant called has reads supporting multiple different variants at the same loci. In the example shown, there were calls for all four nucleotides (A, C, G and T) at the same loci, making it an unreliable call and unlikely to be a true somatic variant.

*Helpful Hints:*

1) Make sure you scroll all the way to the bottom of the track to visualize all of the reads. It is not enough just to rely on the coverage coloring as there might be multiple variants that have a VAF too small to be represented in the coverage bar.

2) Clicking the coverage track will give you the relative abundance of each base at the selected loci.

3) For very deep data, multiple variants due to random error will start to accumulate. The relative abundance, or variant allele frequency, of each base should be considered in cases with deep coverage.

**Figure S14.**
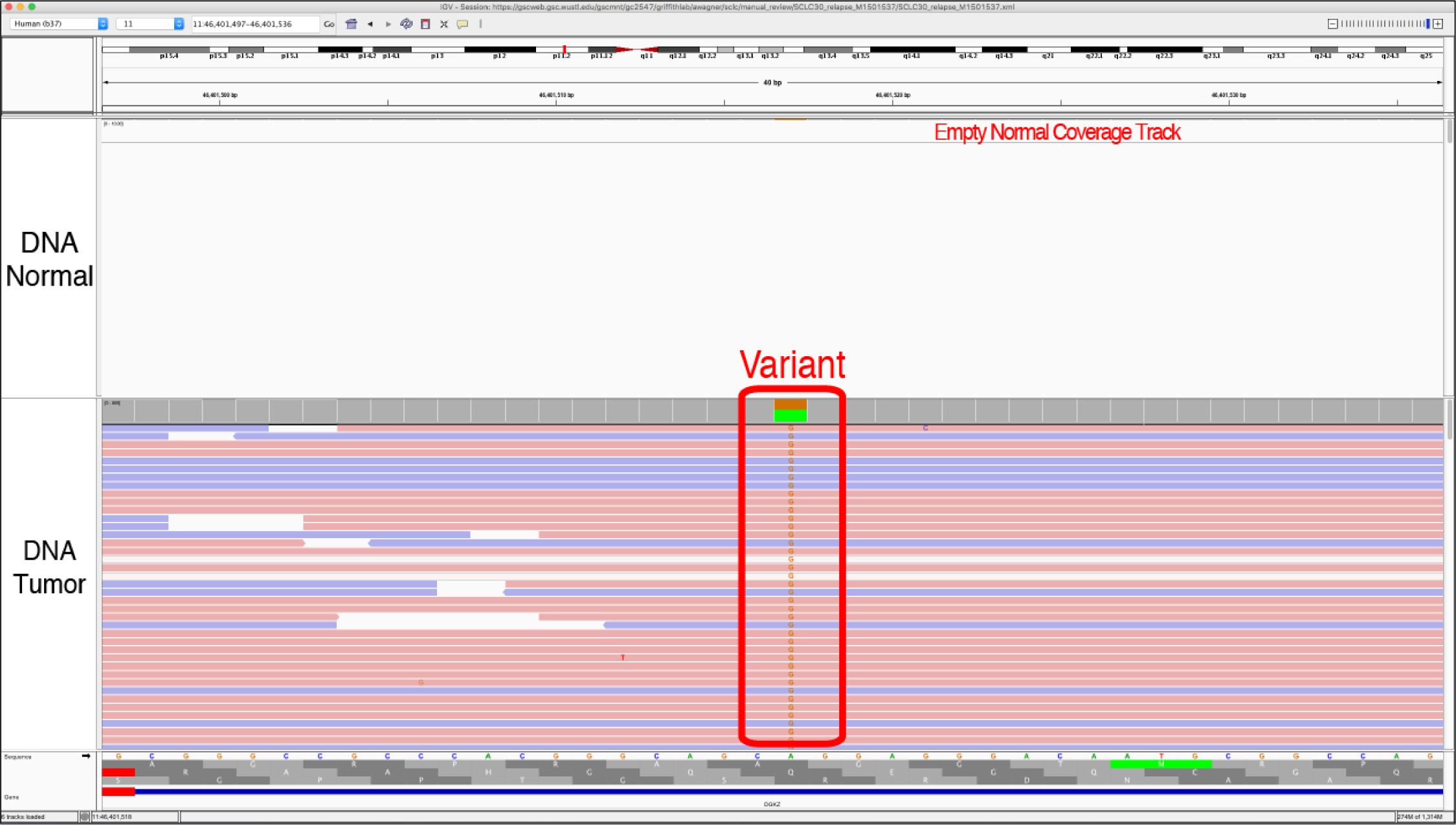
Example of No Coverage in Normal (NCN). The NCN tag is used when there is no coverage in the normal sample with which to effectively compare the tumor sample. This can occur when you do not have a normal track for comparison or you do not have any reads in the normal track to help assess the variant. Typically, at least 20X coverage in both tracks is required to be sure that a call is truly somatic.

*Helpful Hints:*

1) No Count Normal will always occur when you are evaluating tumor only samples, however, the tag is typically used when there should be coverage in the normal track but for a specific variant the coverage is absent.

2) Thresholds can be used pre-filter variants with low VAF in tumor or normal to eliminate the need to evaluate these variants during manual review.

**Figure S15.**
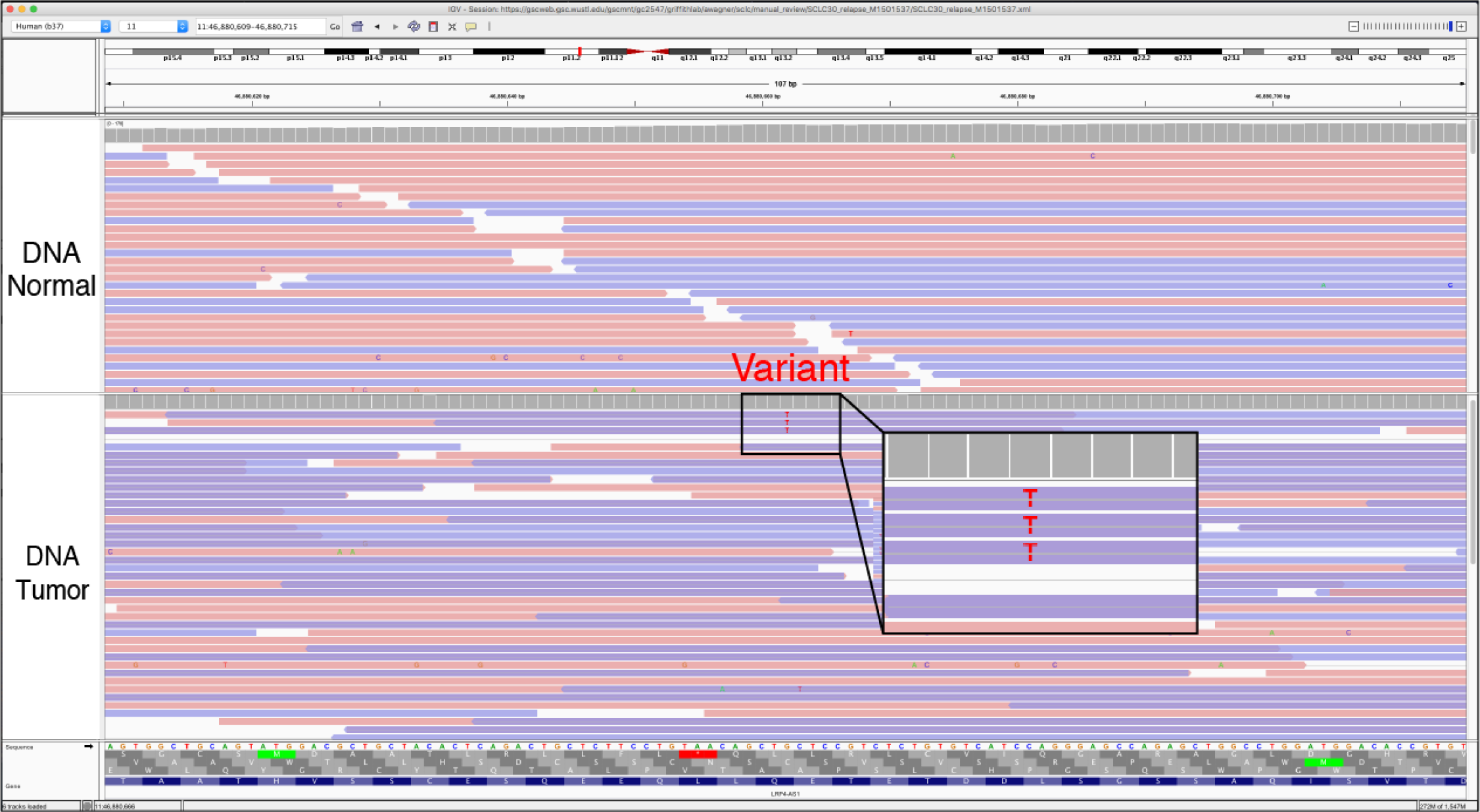
Example of Short Inserts (SI) and Short Inserts Only (SIO). Short inserts refer to instances when the DNA fragment is small enough that sequencing from each end of the molecule results in overlapping reads. That is, the variant appears in the overlapping region of the two read fragments, indicated by the line through the middle of the reads (when viewing reads as pairs). Variants supported by read pairs produced from these short fragments can result in the appearance of two independent reads supporting a variant when in reality, they represent only a single molecule of DNA. The SI tag is used when support for the called variant is primarily from short-insert read pairs but other read strands that show variant support are not short inserts. The SIO tag is used when support for the called variant is exclusively present in paired reads from short inserts. This issue is prevalent in data derived from archival material (FFPE) or other source material with small DNA fragments (e.g., cell-free DNA). Artifacts resulting from short inserts are generally observed at lower frequencies, and are present in two or three read pairs (four to six reads in total). Many variant callers do not correctly account for the non-independence of overlapping read support from short inserts because they were developed at a time when read lengths were shorter and this issue had not yet manifested itself.

*Helpful Hints:*

1) To visualize short insert variants you must view the tracks as pairs. When viewing the reads as pairs, the short inserts will be condensed and a grey line in the middle of a read strand will indicate overlap.

2) When the alignments are colored by read strand, short inserts will appear as dark purple bars. At either end of the short insert, the read strand will change colors to appear either blue or pink, which represents areas of non-overlap.

3) Be sure to visualize each variant using the view as pairs feature as well as using the default settings. The view as pairs feature will allow one to visualize the short inserts, however it will also collapse reads to reduce the total information available to the reviewer. For example, condensed reads might make the variant appear to only be only on reads of a specific direction or it might make the supporting reads appear to be at the ends of read strands. Toggling the feature will prevent reviewers from assigning incorrect tags.

**Figure S16.**
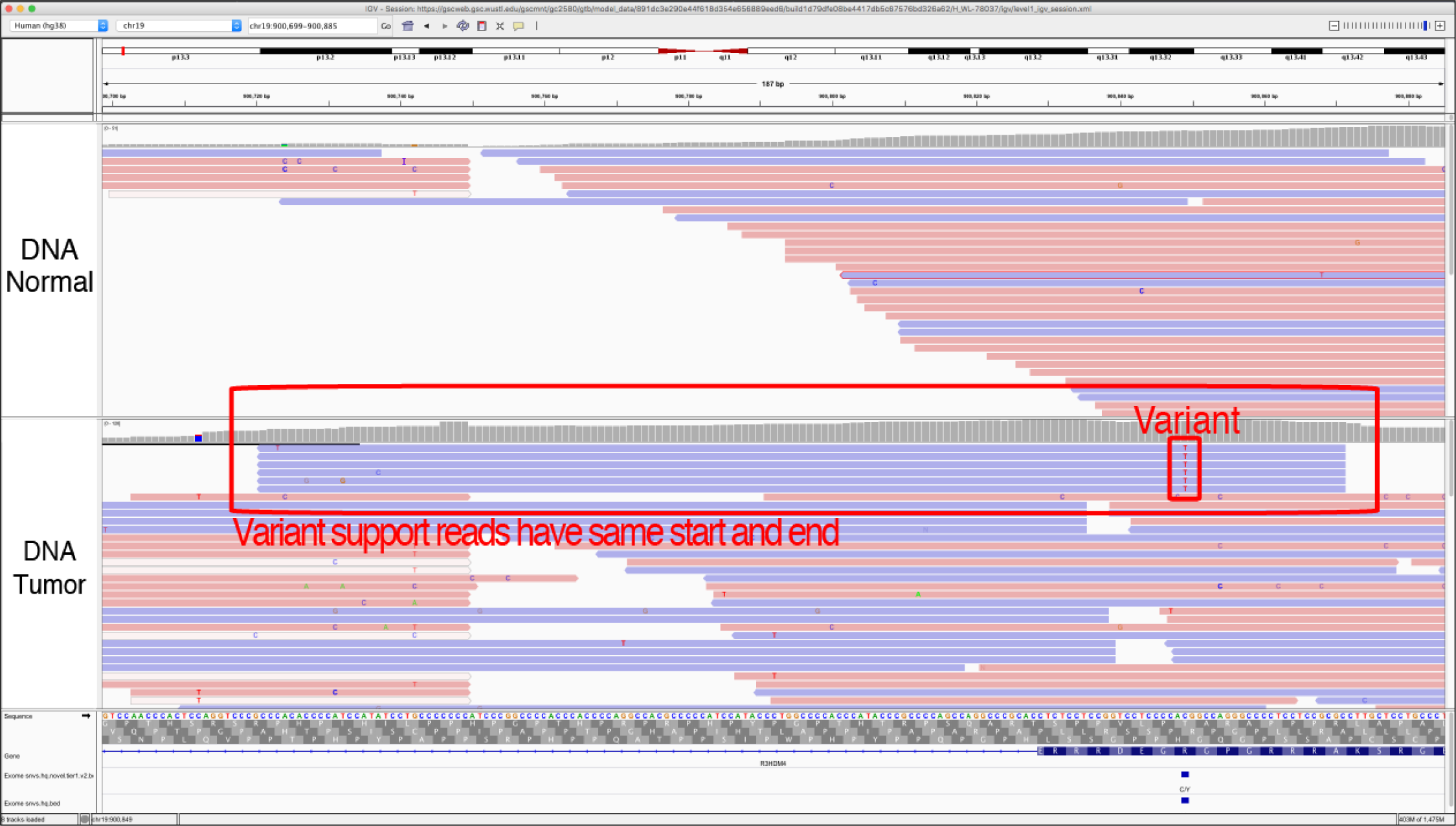
Example of Same Start/End (SSE). The Same Start/End pattern occurs when the variant is only contained by reads that start and stop at the same loci. This is typically attributed to a variant called in multiple reads created from the same originating molecule during the library amplification process, but erroneously not removed during read deduplication.

*Helpful Hints:*

1) Make sure you sort by the variant and zoom out to show the entire length of the reads. This will allow you to visualize if the read stars and ends are the same.

**Figure S17.**
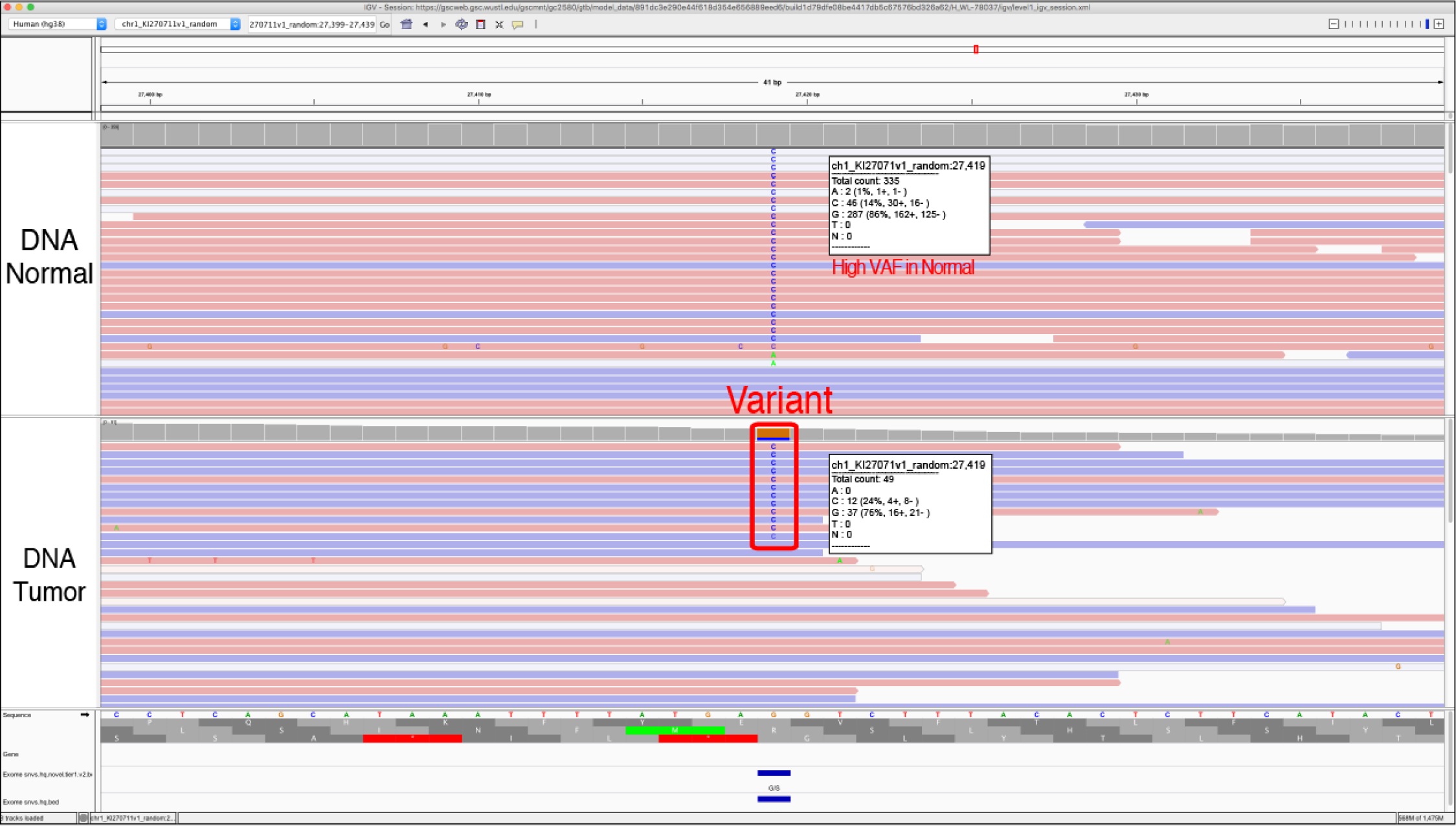
Example of Tumor in Normal (TN). The tumor in normal (TN) is used to indicate that the variant has reads of support in the normal track(s). This is a common occurrence in blood tumors (leukemia and lymphoma) as well as tumors that are highly metastatic. Although this might not be a reason for failing the variant call, it can be used in cases of ambiguity to denote reasons for potential failure. Variants created by sequencing or alignment artifacts will also often occur in both the tumor and the normal sample.

*Helpful Hints:*

1) This does not occur in all hematopoietic tumors but is likely when tumor cells are circulating in the blood stream such as acute myeloid leukemias with high blast counts.

2) Similarly, tumors that are metastatic may have tumor cells circulating in the blood stream and thus can also have tumor in normal contamination.

3) Evaluating other normal samples from your cohort, or evaluating multiple variants within the same sample/experiment, can help set a relative threshold for allowable tumor in normal. This will help to differentiate sequencing and pipeline artifacts from tumor contamination of normal tracks.

**Figure S18.**
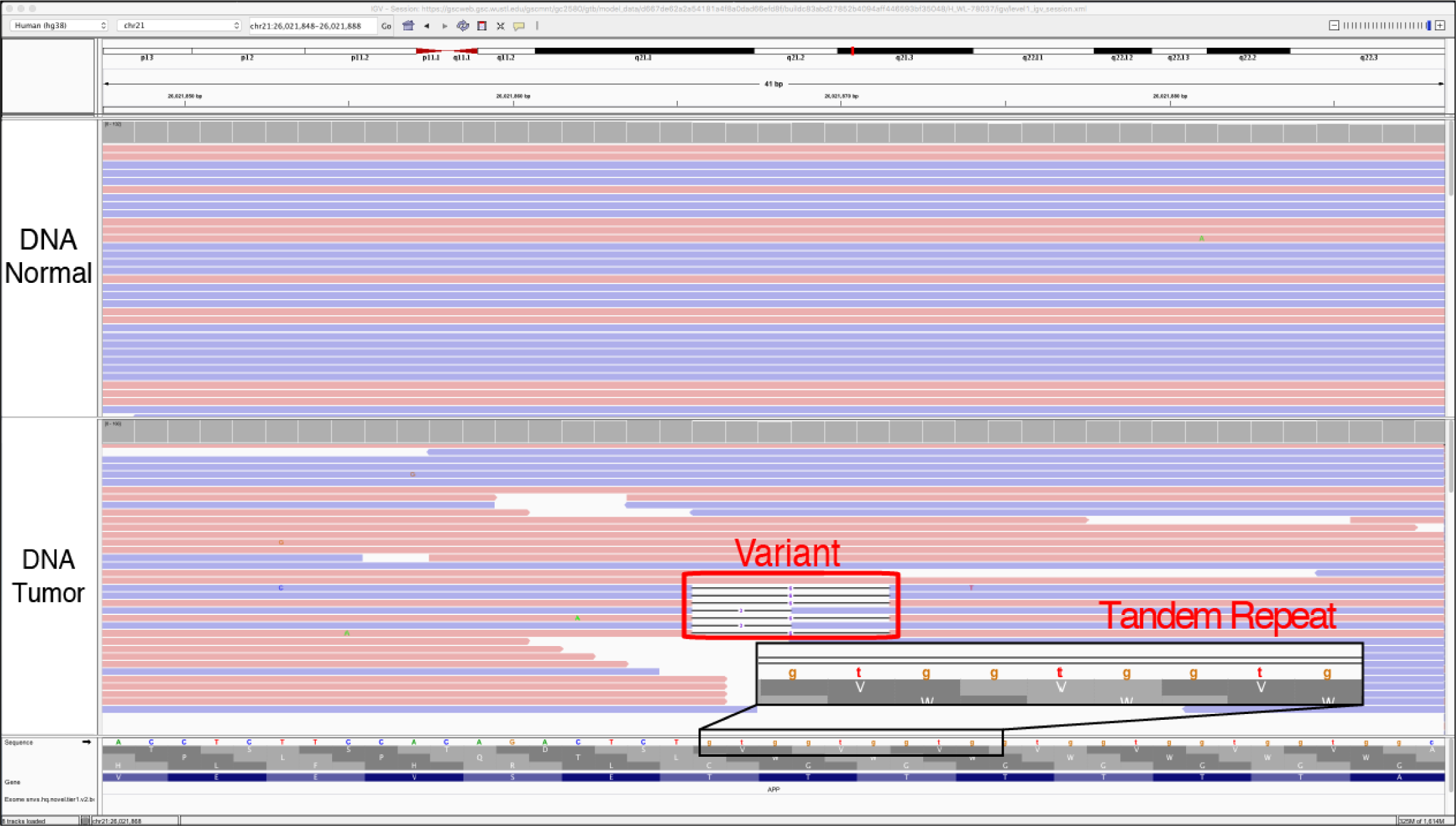
Example of Tandem Repeat (TR). The tandem repeat artifact refers to when the reference sequence contains a repeat region of some number of nucleotides (e.g. GTGGTGGTG…). The variant being evaluated may be a SNV or indel directly adjacent or within this region and called due to misalignment or errors at the ends of reads. Repeat regions are areas where some sequencers, particularly those dependent on the polymerase enzyme, are prone to making mistakes. However, it is important to note that these are also areas of normal human variation and real *de novo* mutations due to errors produced by polymerase during DNA replication. Like the Mononucleotide (MN) and Dinucleotide (DN) tags, other factors such as the size of the repeat, appearance in the normal and indels of varying length should be considered during evaluation.

*Helpful Hints:*

1) Typically, these variants are small deletions or small insertions and they are usually visualized in both the tumor tracks and the normal tracks.

2) Although the variant being evaluated may be an n-bp deletion, deletions of different sizes or even insertions are often observed at these sites.

## References

1. Shiraishi Y, Sato Y, Chiba K et al. An empirical Bayesian framework for somatic mutation detection from cancer genome sequencing data. Nucleic Acids Res 2013; 41: e89.

2. Cibulskis K, Lawrence MS, Carter SL et al. Sensitive detection of somatic point mutations in impure and heterogeneous cancer samples. Nat Biotechnol 2013; 31: 213–219.

3. Larson DE, Harris CC, Chen K et al. SomaticSniper: identification of somatic point mutations in whole genome sequencing data. Bioinformatics 2012; 28: 311–317.

4. Saunders CT, Wong WSW, Swamy S, Becq J, Murray LJ, Cheetham RK. Strelka: accurate somatic small-variant calling from sequenced tumor-normal sample pairs. Bioinformatics 2012; 28:1811–1817.

5. Koboldt DC, Zhang Q, Larson DE et al. VarScan 2: somatic mutation and copy number alteration discovery in cancer by exome sequencing. Genome Res 2012; 22: 568–576.

6. Kim S, Jeong K, Bhutani K et al. Virmid: accurate detection of somatic mutations with sample impurity inference. Genome Biol 2013; 14: R90.

7. Krøigård AB, Thomassen M, Lænkholm A-V, Kruse TA, Larsen MJ. Evaluation of Nine Somatic Variant Callers for Detection of Somatic Mutations in Exome and Targeted Deep Sequencing Data. PLoS One 2016; 11: e0151664.

8. Cai L, Yuan W, Zhang Z, He L, Chou K-C. In-depth comparison of somatic point mutation callers based on different tumor next-generation sequencing depth data. Sci Rep 2016; 6: 36540.

9. Callari M, Sammut S-J, De Mattos-Arruda L et al. Intersect-then-combine approach: improving the performance of somatic variant calling in whole exome sequencing data using multiple aligners and callers. Genome Med 2017; 9: 35.

10. Poplin R, Newburger D, Dijamco J et al. Creating a universal SNP and small indel variant caller with deep neural networks. 2016. doi: 10.1101/092890.

11. Robinson JT, Thorvaldsdóttir H, Wenger AM, Zehir A, Mesirov JP. Variant Review with the Integrative Genomics Viewer. Cancer Res 2017; 77: e31–e34.

12. Thorvaldsdottir H, Robinson JT, Mesirov JP. Integrative Genomics Viewer (IGV): high-performance genomics data visualization and exploration. Brief Bioinform 2012; 14: 178–192.

13. Strom SP. Current practices and guidelines for clinical next-generation sequencing oncology testing. Cancer Biol Med 2016; 13: 3–11.

14. Sukhai MA, Craddock KJ, Thomas M et al. A classification system for clinical relevance of somatic variants identified in molecular profiling of cancer. Genet Med 2016; 18: 128–136.

15. Kim J, Park W-Y, Kim NKD et al. Good Laboratory Standards for Clinical Next-Generation Sequencing Cancer Panel Tests. J Pathol Transl Med 2017; 51: 191–204.

16. Rasche L, Chavan SS, Stephens OW et al. Spatial genomic heterogeneity in multiple myeloma revealed by multi-region sequencing. Nat Commun 2017; 8: 268.

17. Ott PA, Hu Z, Keskin DB et al. An immunogenic personal neoantigen vaccine for patients with melanoma. Nature 2017; 547: 217–221.

18. Rheinbay E, Parasuraman P, Grimsby J et al. Recurrent and functional regulatory mutations in breast cancer. Nature 2017; 547: 55–60.

19. Giannakis M, Hodis E, Mu XJ et al. RNF43 is frequently mutated in colorectal and endometrial cancers. Nat Genet 2014; 46: 1264–1266.

20. Robinson DR, Wu Y-M, Lonigro RJ et al. Integrative clinical genomics of metastatic cancer. Nature 2017; 548: 297–303.

21. Fiume M, Williams V, Brook A, Brudno M. Savant: genome browser for high-throughput sequencing data. Bioinformatics 2010; 26: 1938–1944.

22. Goecks J, Coraor N, Galaxy Team, Nekrutenko A, Taylor J. NGS analyses by visualization with Trackster. Nat Biotechnol 2012; 30: 1036–1039.

23. Carver T, Harris SR, Otto TD, Berriman M, Parkhill J, McQuillan JA. BamView: visualizing and interpretation of next-generation sequencing read alignments. Brief Bioinform 2013; 14:203–212.

24. Griffith M, Griffith OL, Smith SM et al. Genome Modeling System: A Knowledge Management Platform for Genomics. PLoS Comput Biol 2015; 11: e1004274.

25. 1000 Genomes Project Consortium, Auton A, Brooks LD et al. A global reference for human genetic variation. Nature 2015; 526: 68–74.

26. Lek M, Karczewski KJ, Minikel EV et al. Analysis of protein-coding genetic variation in 60,706 humans. Nature 2016; 536: 285–291.

27. Yost SE, Smith EN, Schwab RB et al. Identification of high-confidence somatic mutations in whole genome sequence of formalin-fixed breast cancer specimens. Nucleic Acids Res 2012; 40: e107.

28. Akbari M, Hansen MD, Halgunset J, Skorpen F, Krokan HE. Low Copy Number DNA Template Can Render Polymerase Chain Reaction Error Prone in a Sequence-Dependent Manner. J Mol Diagn 2005; 7: 36–39.

29. Walsh PS, Erlich HA, Higuchi R. Preferential PCR amplification of alleles: mechanisms and solutions. PCR Methods Appl 1992; 1:241–250.

30. Aird D, Ross MG, Chen W-S et al. Analyzing and minimizing PCR amplification bias in Illumina sequencing libraries. Genome Biol 2011; 12: R18.

31. Nakamura K, Oshima T, Morimoto T et al. Sequence-specific error profile of Illumina sequencers. Nucleic Acids Res 2011; 39: e90.

32. Griffith M, Spies NC, Krysiak K et al. CIViC is a community knowledgebase for expert crowdsourcing the clinical interpretation of variants in cancer. Nat Genet 2017; 49: 170–174.

